# Fractal signatures of SARS-CoV2 coronavirus, the indicator matrix, the fractal dimension and the 2D directional wavelet transform: A comparative study with SARS-CoV, MERS-CoV and SARS-like coronavirus

**DOI:** 10.1101/2020.08.26.269118

**Authors:** Sid-Ali Ouadfeul

## Abstract

The main goal of this paper is to show the 2D fractal signatures of SARS-CoV2 coronavirus, indicator matrixes maps showing the concentration of nucleotide acids are built form the RNA sequences, and then the fractal dimension and 2D Directional Wavelet Transform (DCWT) are calculated. Analysis of 21 RNA sequences downloaded from NCBI database shows that indicator matrixes and 2D DCWT exhibit the same patterns with different positions, while the fractal dimensions are oscillating around 1.60. A comparison with SARS-CoV, MERS-CoV and SARS-like Coronavirus shows slightly different fractal dimensions, however the indicator matrix and 2D DCWT exhibit the same patterns for the couple (SARS-CoV2, SARS-CoV) and (MERS-CoV, SARS-like) Coronavirus. Obtained results show that SARS-CoV2 is probably a result of SARS-CoV mutation process.

## 1. Introduction

Fractal analysis of SARS-CoV2 Coronavirus genomes is one of the useful tools to characterize the virus, Ouadfeul (2020) published a paper deals with the multifractal analysis of the Coronavirus gnomes using the wavelet transform, six DNA coding methods are used and the Long-Range correlation is investigated. Obtained results show that the SRAS-CoV2 is undergoes mutation, unstable, far from the equilibrium. Fernandes et al (2020) published a paper deals with the investigation of SARS-CoV, MERS-CoV, and SARS-CoV-2 paradigm of chaos theory and fractal geometry. Hassan et al (2020) published a paper deals with the quantitative understanding of the purine and pyrimidine spatial distribution/organization of all 89 complete sequences of SARS-CoV (available as on date in the NCBI virus database), is made using different parameters such as fractal dimension, Hurst exponent, Shannon entropy and GC content of the nucleotide sequences of the genome of SARS-CoV2. Also a cluster among all the the SARS-CoV sequences of nucleotide have been made based on their phylogeny made through their closeness (Hamming distance) based on respective purine-pyrimidine distribution. Mandal et al (2020) published studied and analyzed a large number of publicly available SARS-CoV-2 genomes across the world using the multifractal approach. The mutation events in the isolates obey the Markov process and exhibit very high mutational rates. In this paper, the indicator matrix of 21 RNA sequences downloaded from the NCBI database, and then the fractal dimension and 2D directional wavelet transform (DCWT) are calculated, a comparison with MERS-CoV, SARS-CoV and SARS-like coronavirus is done. We begin the paper by talking about the indicator matrix and the fractal dimension.

## 2. The indicator matrix and the fractal dimension

The DNA of each organism of a given species is a long sequence of a specific large number of base pairs bp. Each base pair is defined on the 4 elements alphabet of nucleotides (Cattani, 2010):

A: adenine, T: thymine, C: cytosine, G; guanine

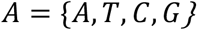

A DNA sequence is the finite symbolic sequence

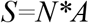

*S* is defined as:

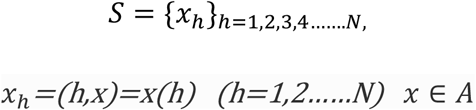

***x_h_*** is the value of *x* at the position *h*

The 2D indicator function, based on the 1D-definition is the map:

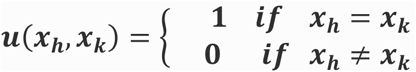

The indicator matrix *C* is defined as:

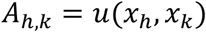

*A* is a square matrix with dimension *N*N*

Table 1 shows an example of construction of the indicator matrix.

**Table 1:**
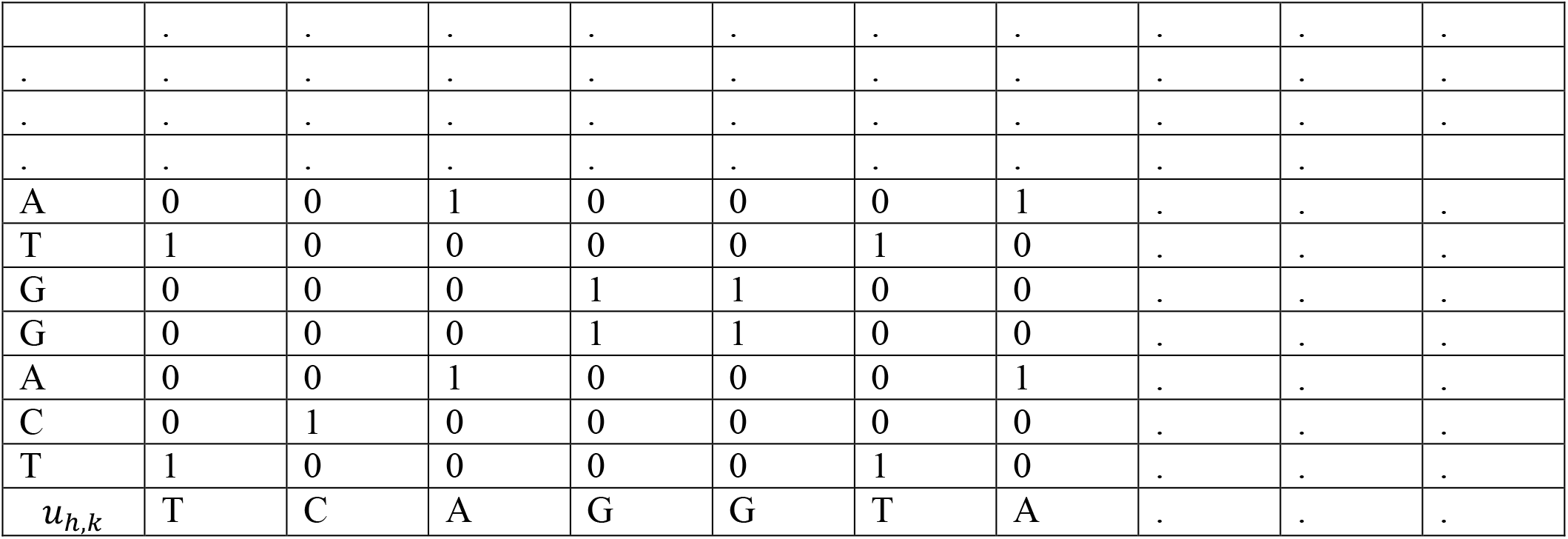
The indicator matrix components

|  |  |  |  |  |  |  |  |  |  |  |
| --- | --- | --- | --- | --- | --- | --- | --- | --- | --- | --- |
|  | . | . | . | . | . | . | . | . | . | . |
| . | . | . | . | . | . | . | . | . | . | . |
| . | . | . | . | . | . | . | . | . | . | . |
| . | . | . | . | . | . | . | . | . | . | . |
| A | 0 | 0 | 1 | 0 | 0 | 0 | 1 | . | . | . |
| T | 1 | 0 | 0 | 0 | 0 | 1 | 0 | . | . | . |
| G | 0 | 0 | 0 | 1 | 1 | 0 | 0 | . | . | . |
| G | 0 | 0 | 0 | 1 | 1 | 0 | 0 | . | . | . |
| A | 0 | 0 | 1 | 0 | 0 | 0 | 1 | . | . | . |
| C | 0 | 1 | 0 | 0 | 0 | 0 | 0 | . | . | . |
| T | 1 | 0 | 0 | 0 | 0 | 1 | 0 | . | . | . |
| $u_{h,k}$ | T | C | A | G | G | T | A | . | . | . |

From the indicator matrix we can have an idea of the “fractal-like” distribution of nucleotides. The fractal dimension for the graphical representation of the indicator matrix plots can be computed as the average of the number p(n) of “1” in the randomly taken *nxn* minors of the *NxN* correlation matrix *u_h,k_* (Cattani, 2010) :

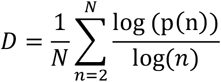

## 3. The 2D Directional Continuous Wavelet Transform

The 2D Directional Continuous Wavelet Transform (DCWT) was introduced by Murenzi (1989). The wavelet decomposition of a given function *f* ∈ *L*^2^(*R*^2^) with an analyzing wavelet *g* ∈ *L*^2^(*R*^2^) is defined for all a>0, *b* ∈ *R*^2^, *α* ∈ [0,2*π*], by :

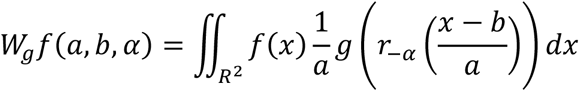

Where *r_−α_* is a rotation with an angle −*α* in the *R*^2^.

An example of a directional wavelet transform is the gradient of the Gaussian Wavelet ∇*G*, since the convolution of an image with ∇*G* is equivalent to analysis of the gradient of the modulus of the continuous wavelet transform. Canny has introduced another tool for edge detection (Arneodo et al, 2003).After the convolution of the image with a Gaussian wavelet, we calculate its gradient to seek the set of points corresponding to maxima locations in the intensity of the 2D continuous wavelet transform. The use of the gradient of the Gaussian as an analyzing wavelet was introduced by Mallat and Hwang (1992).

The wavelet transform of a function f, with an analyzing wavelet *g* = ∇*G* is a vector defined for all a>0, *b* ∈ *R*^2^ defined by:

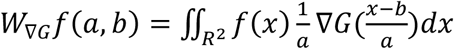

If we choose 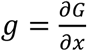, this transformation relates the directional wavelet transform by the following relation:

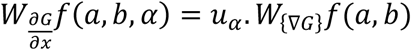

Where *u_α_* is the unit vector in the directions: *u_α_*(cos(*α*), sin(*α*)), and “.” is the Euclidian scalar product in *R*^2^.

In this paper, we choose the Mexican Hat as an analyzing wavelet:

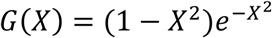

## 4. Application to SARS-CoV2

The indicator matrix of 21 RNA sequences of SARS-CoV2 coronavirus downloaded from NCBI database are calculated, table 02 shows the code of each sequence and the origin of each infected patient. Figure 01 shows the indicator matrix of each sequence, we observe some same patterns in the 21 sequences, and these patterns are changing position for a matrix to another. The fractal dimensions of these indicator matrixes are calculated, table 02 shows the fractal dimension of each sequence. We can observe that these fractal dimensions are oscillating around 1.63 (*D* = 1.63 ± 0.03).

**Table 2.** Fractal dimension of 21 RNA sequences of SARS-CoV coronavirus

| Seq Num | GenBank | Country | Fractal Dimension |
| --- | --- | --- | --- |
| 1 | MT066156.1 | Italy | 1.64 |
| 2 | MT044257 | USA IL | 1.64 |
| 3 | LC528232.1 | Japan | 1.63 |
| 4 | LC528233 | Japan | 1.65 |
| 5 | LR757998.1 | China: Wuhan | 1.66 |
| 6 | MN938384.1 | China: Shenzhen | 1.63 |
| 7 | LR757996.1 | China: Wuhan | 1.62 |
| 8 | MN938384.1 | China: Shenzhen | 1.63 |
| 9 | MT135043.1 | China : Beijing | 1.64 |
| 10 | MT126808.1 | Brazil | 1.64 |
| 11 | MT066175.1 | Taiwan | 1.64 |
| 12 | LC529905.1 | Japan | 1.64 |
| 13 | MN985325.1 | USA WA | 1.64 |
| 14 | MT093571.1 | Sweeden | 1.64 |
| 15 | MN994467 | USA : CA | 1.60 |
| 16 | MT072688 | Nepal | 1.60 |
| 17 | MT007544.1 | Australia Victoria | 1.64 |
| 18 | MT012098.1 | India: Kerala State | 1.63 |
| 19 | MT121215.1 | China Changai | 1.60 |
| 20 | MT198652 | Spain : Valencia | 1.61 |
| 21 | MT192773.1 | Viet Nam | 1.64 |

**Figure 01:**
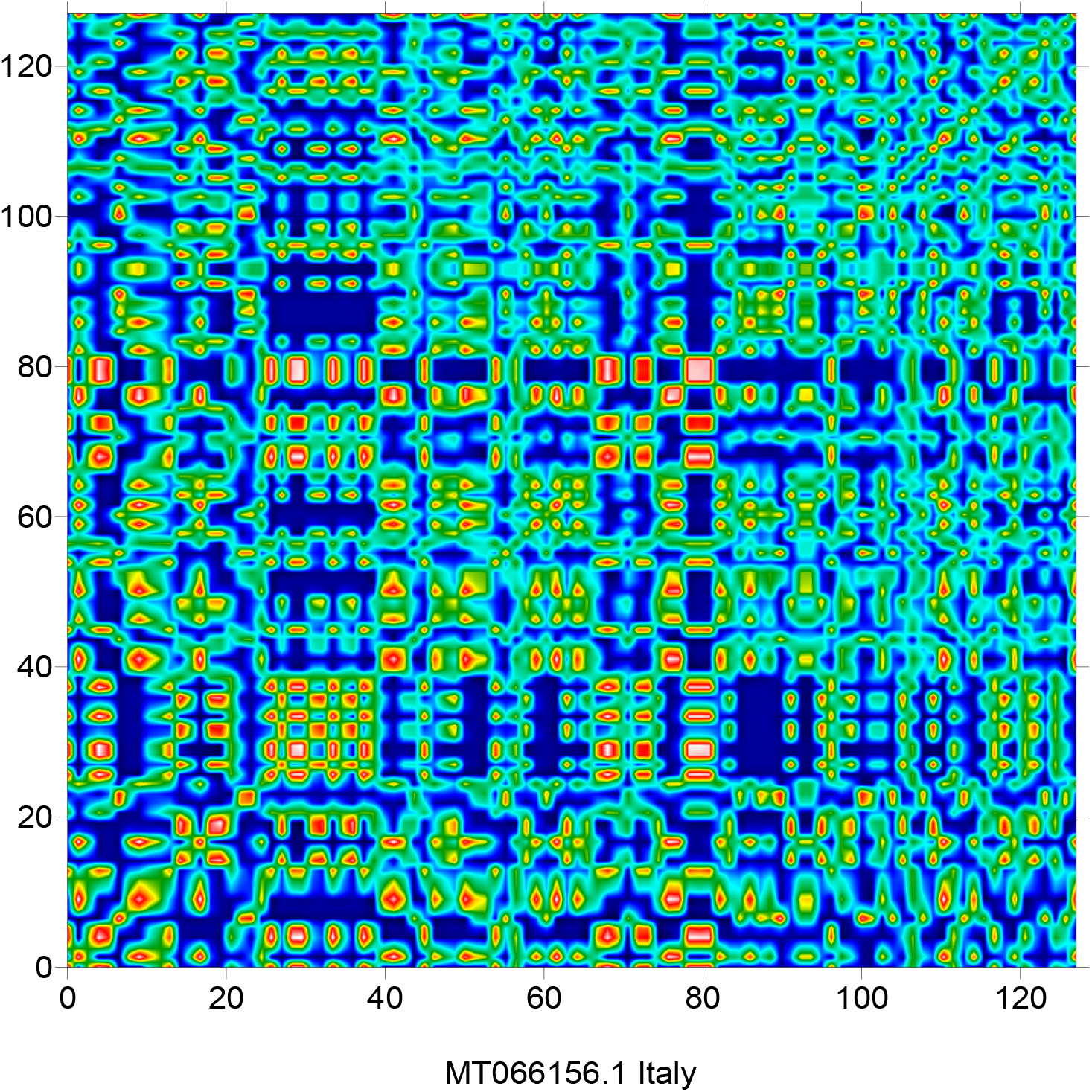
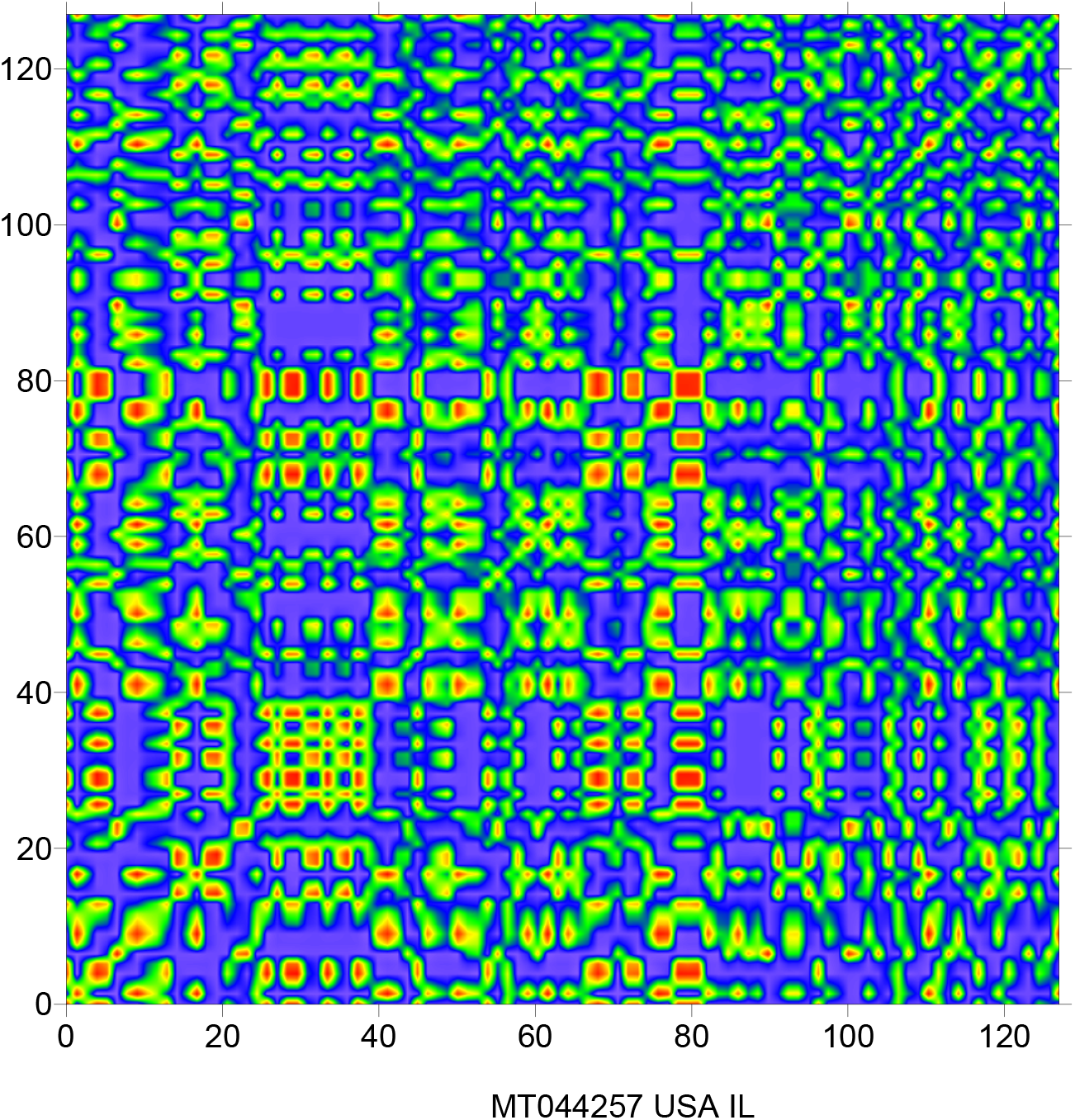
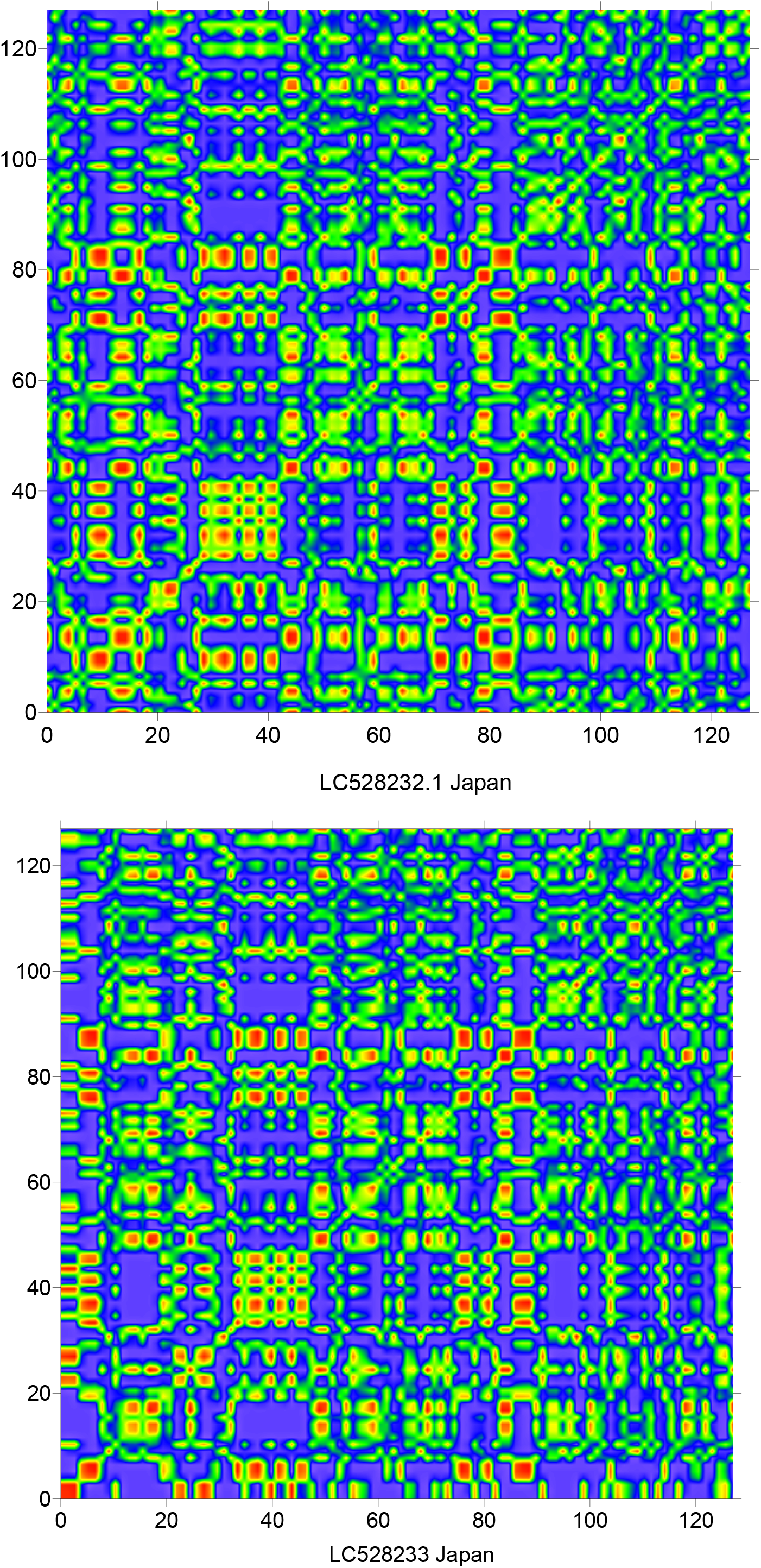
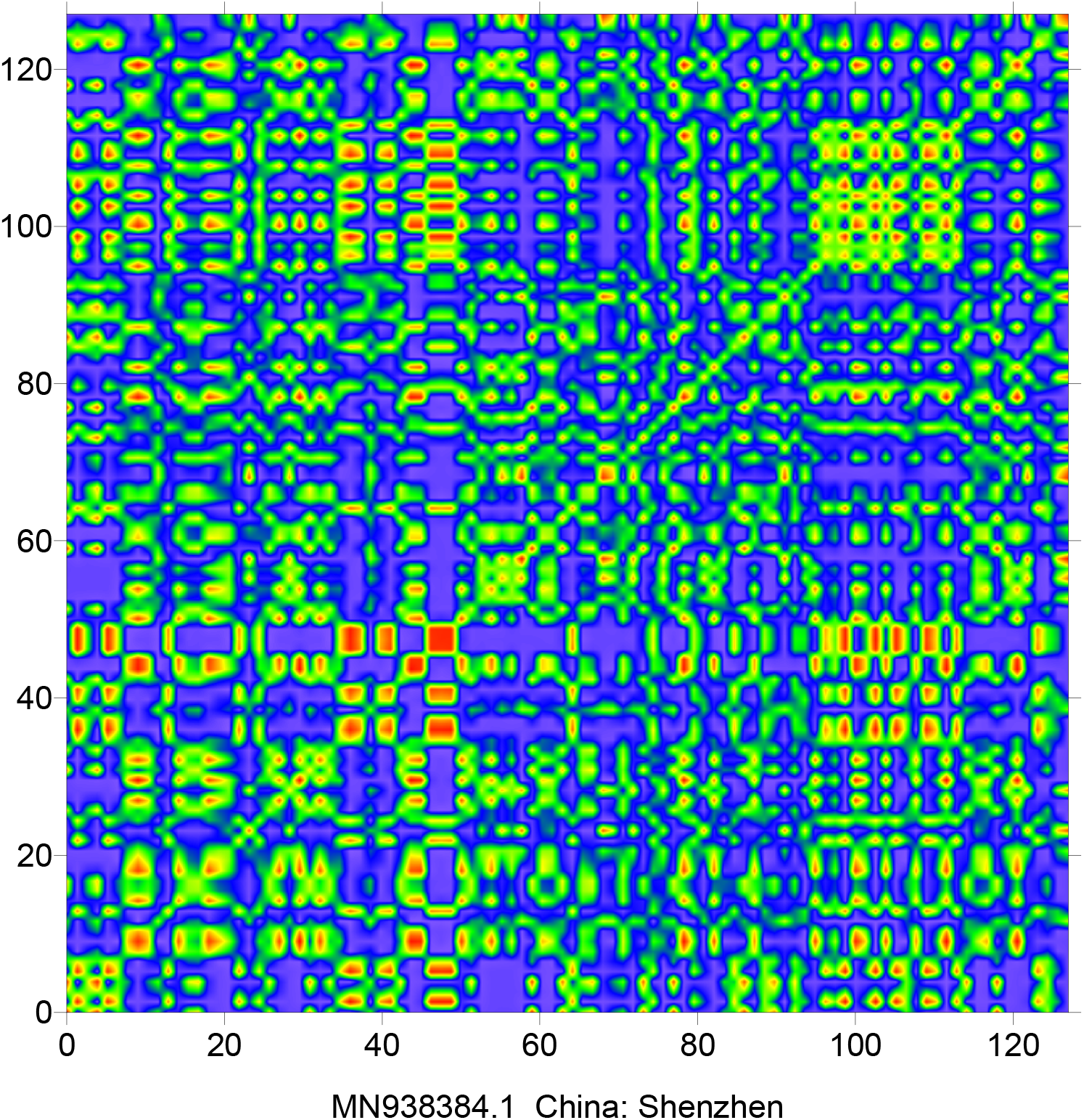
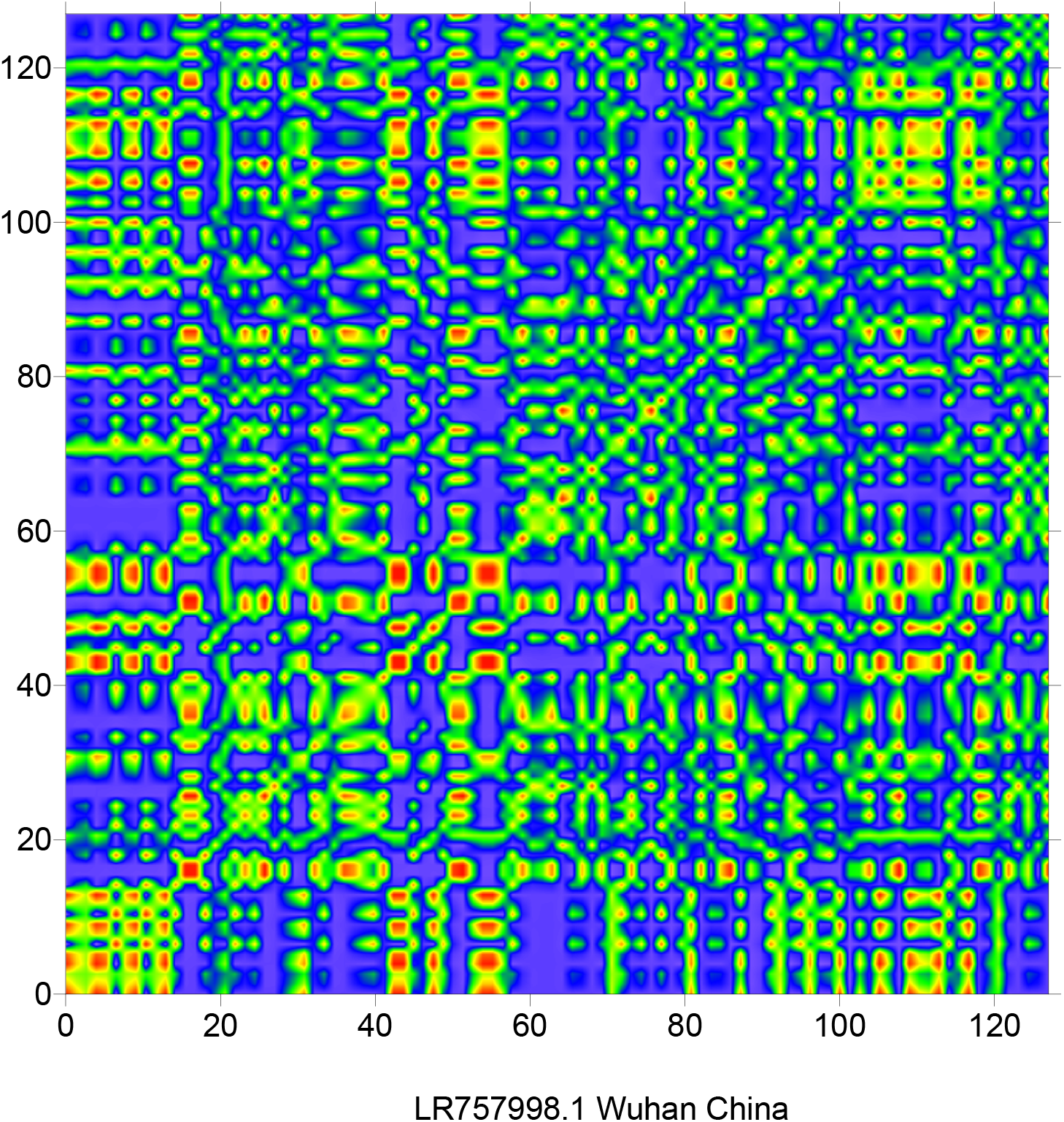
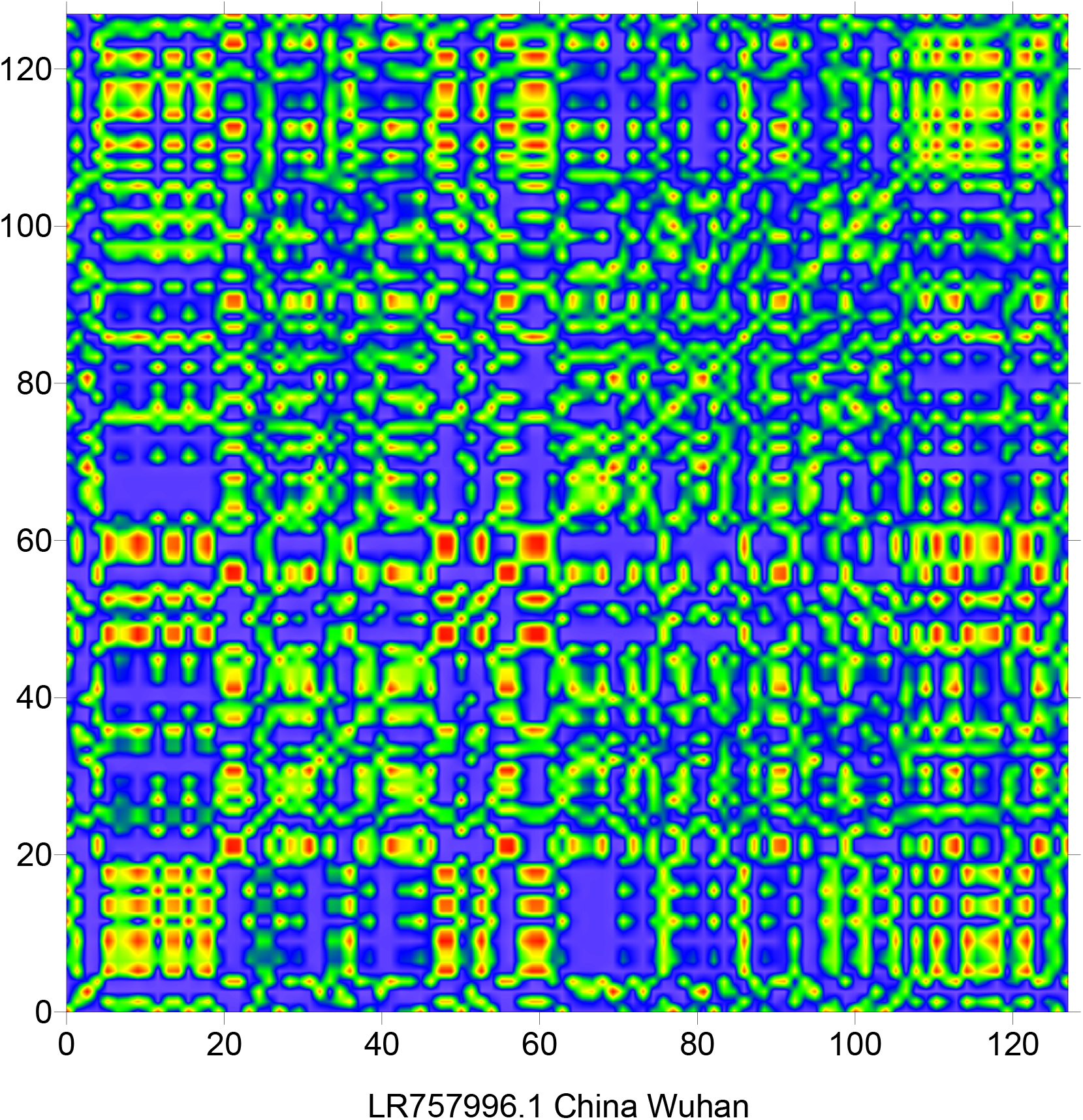
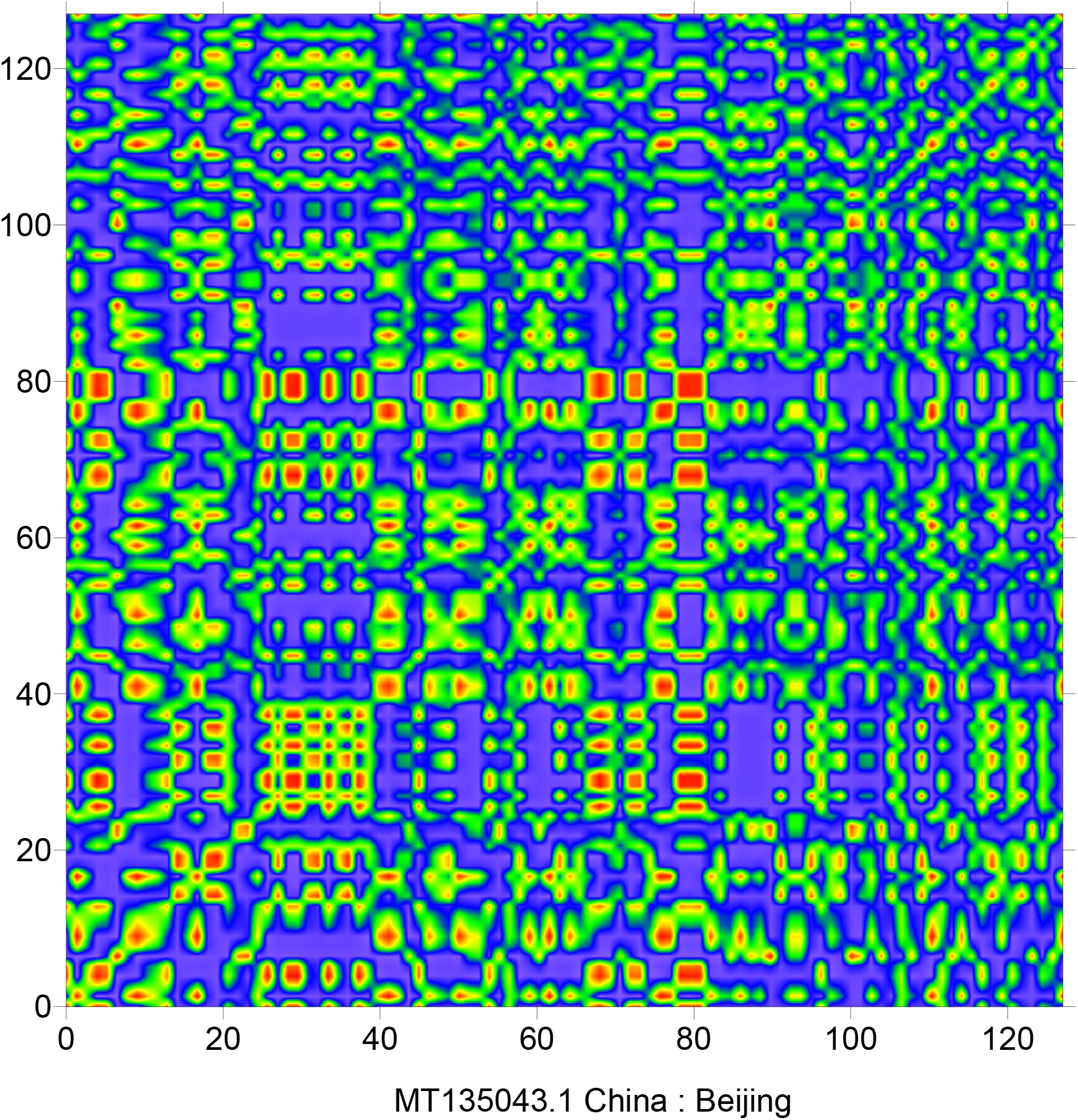
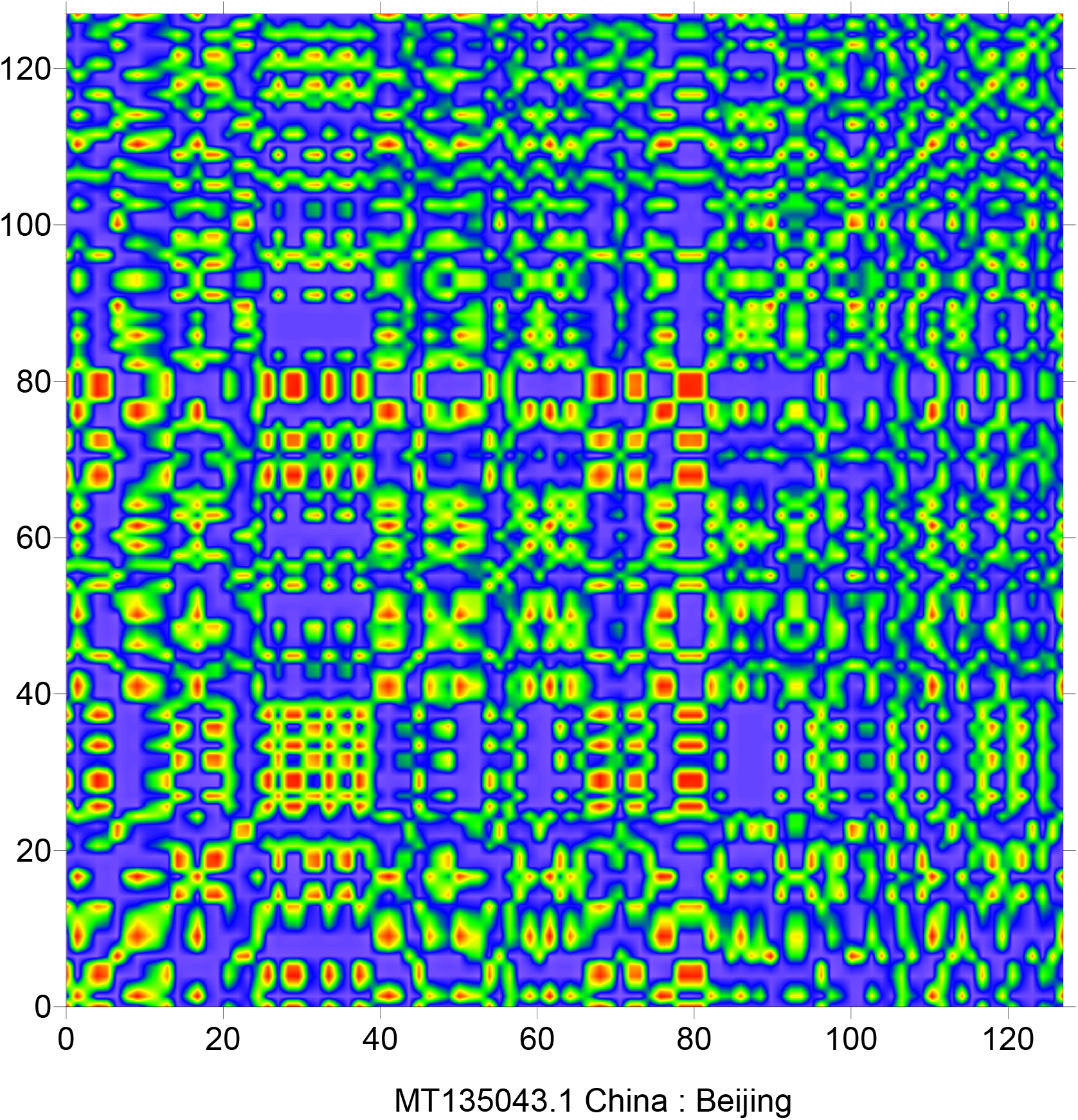
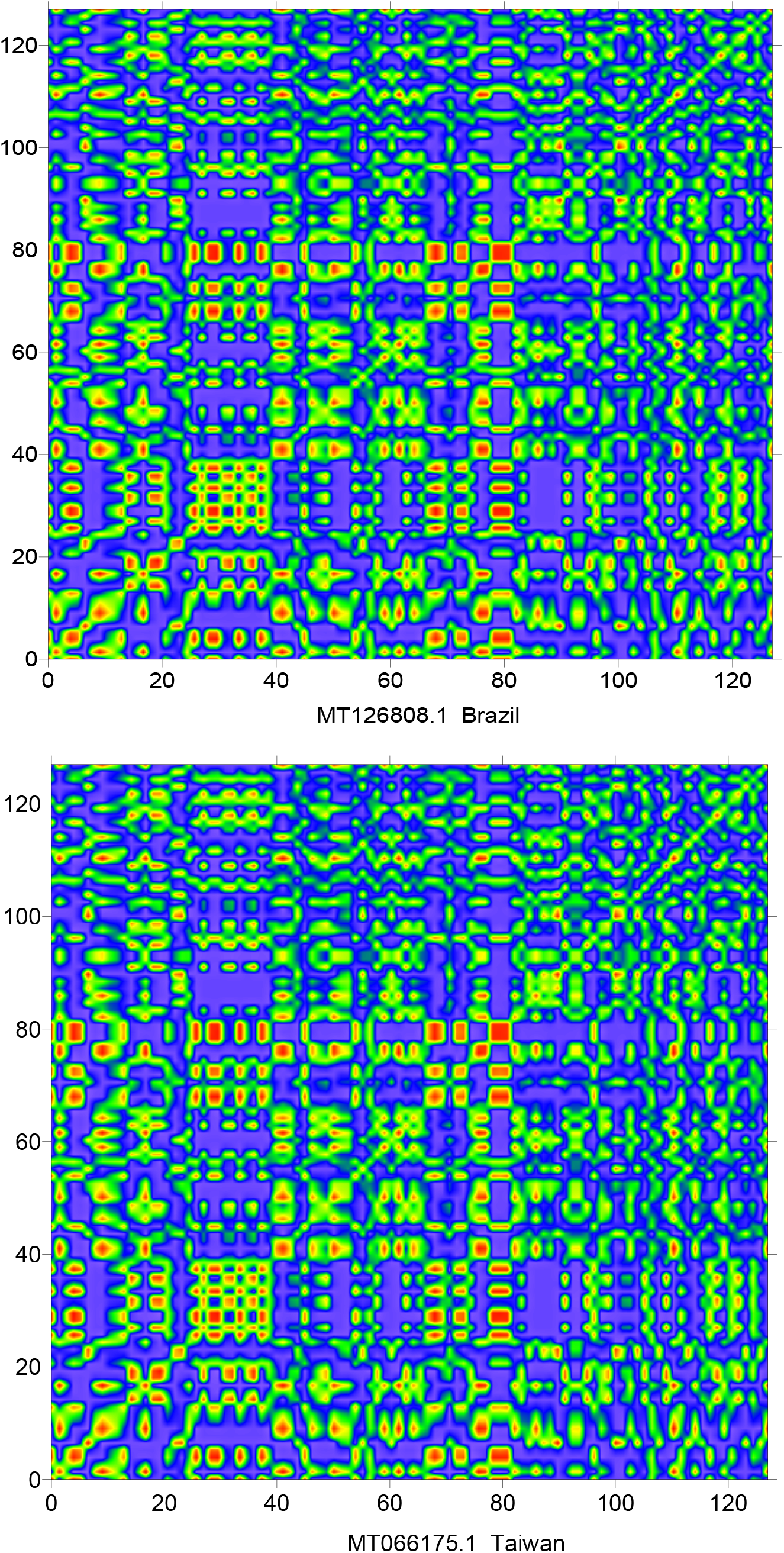
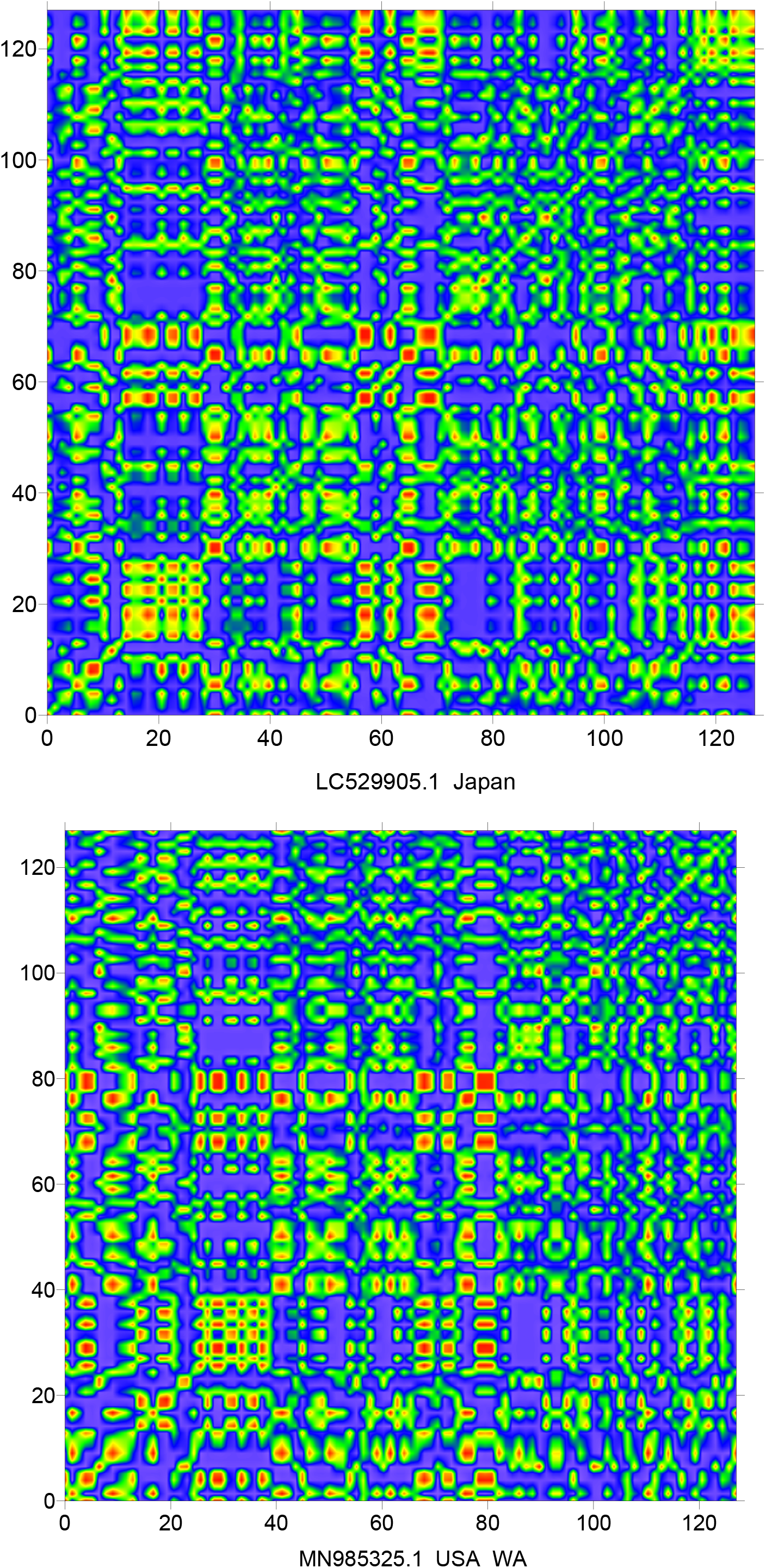
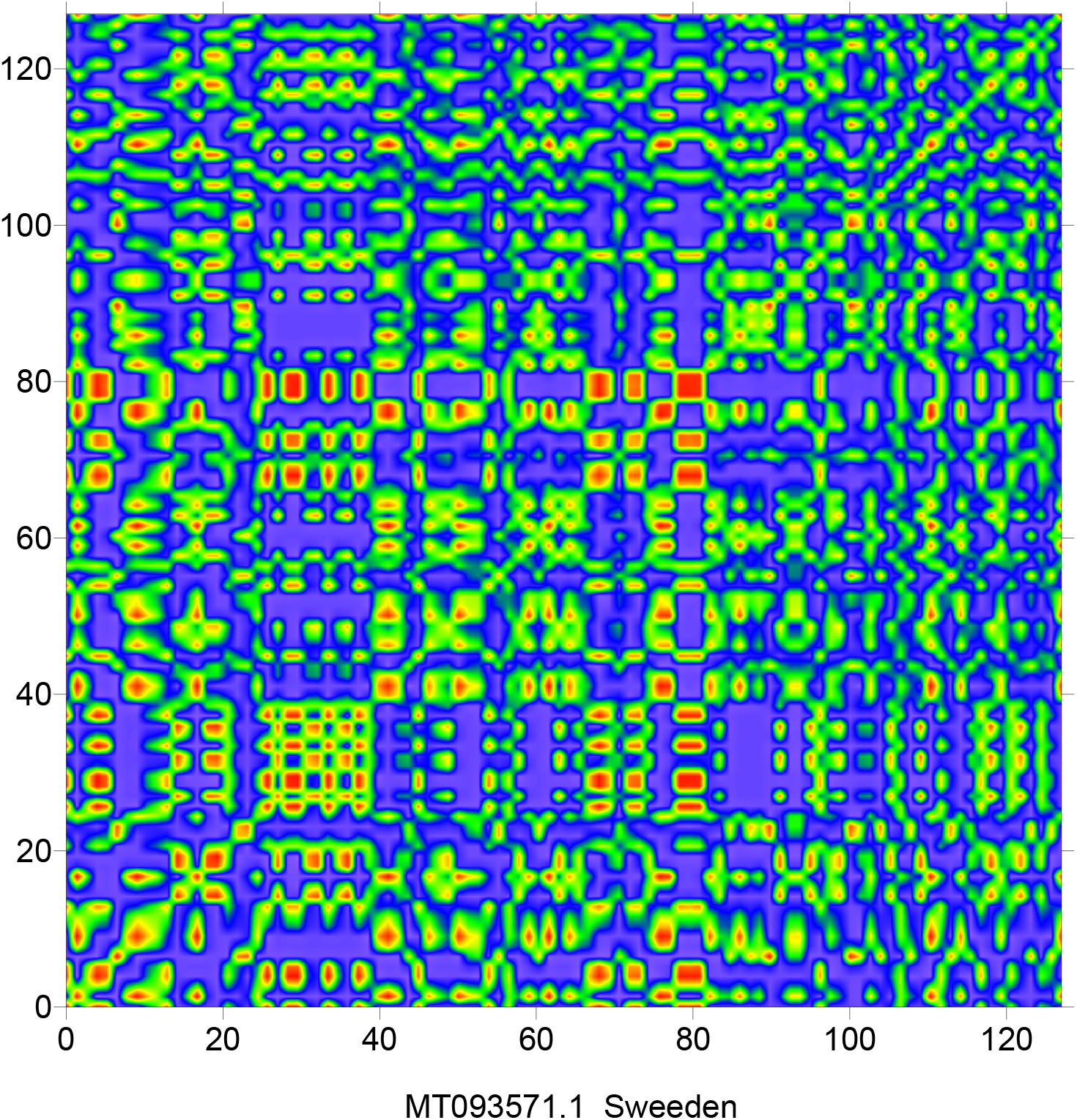
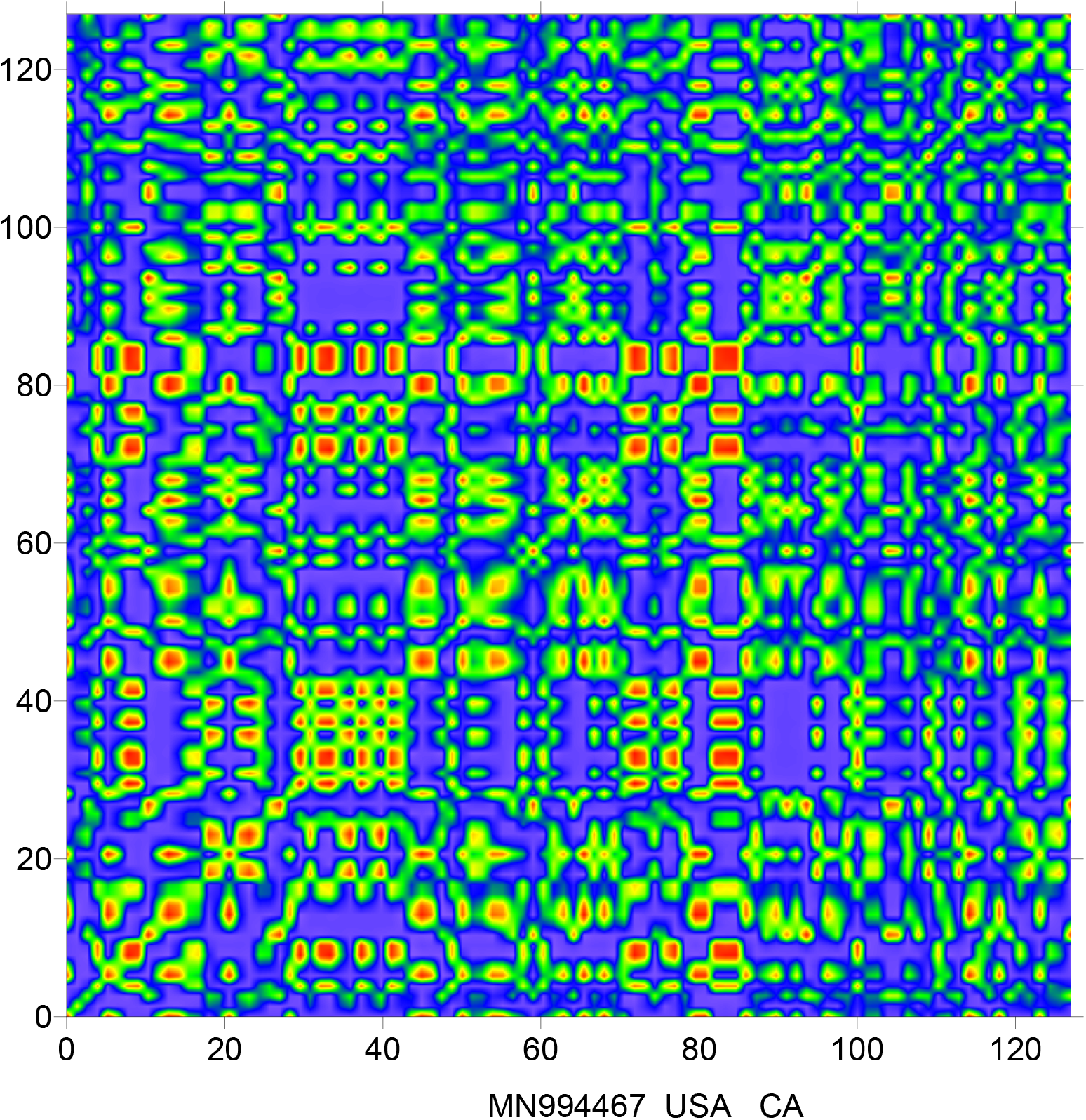
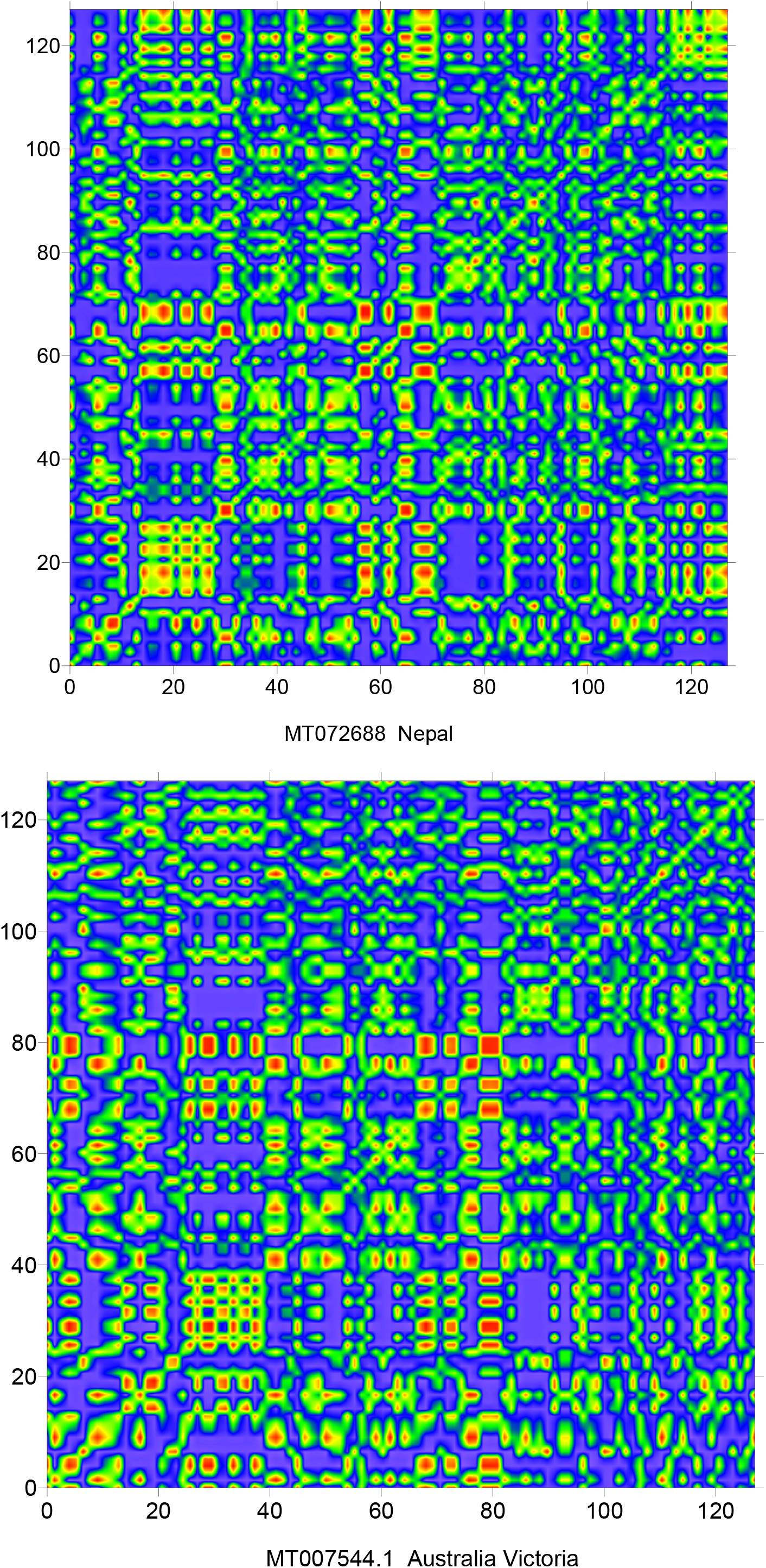
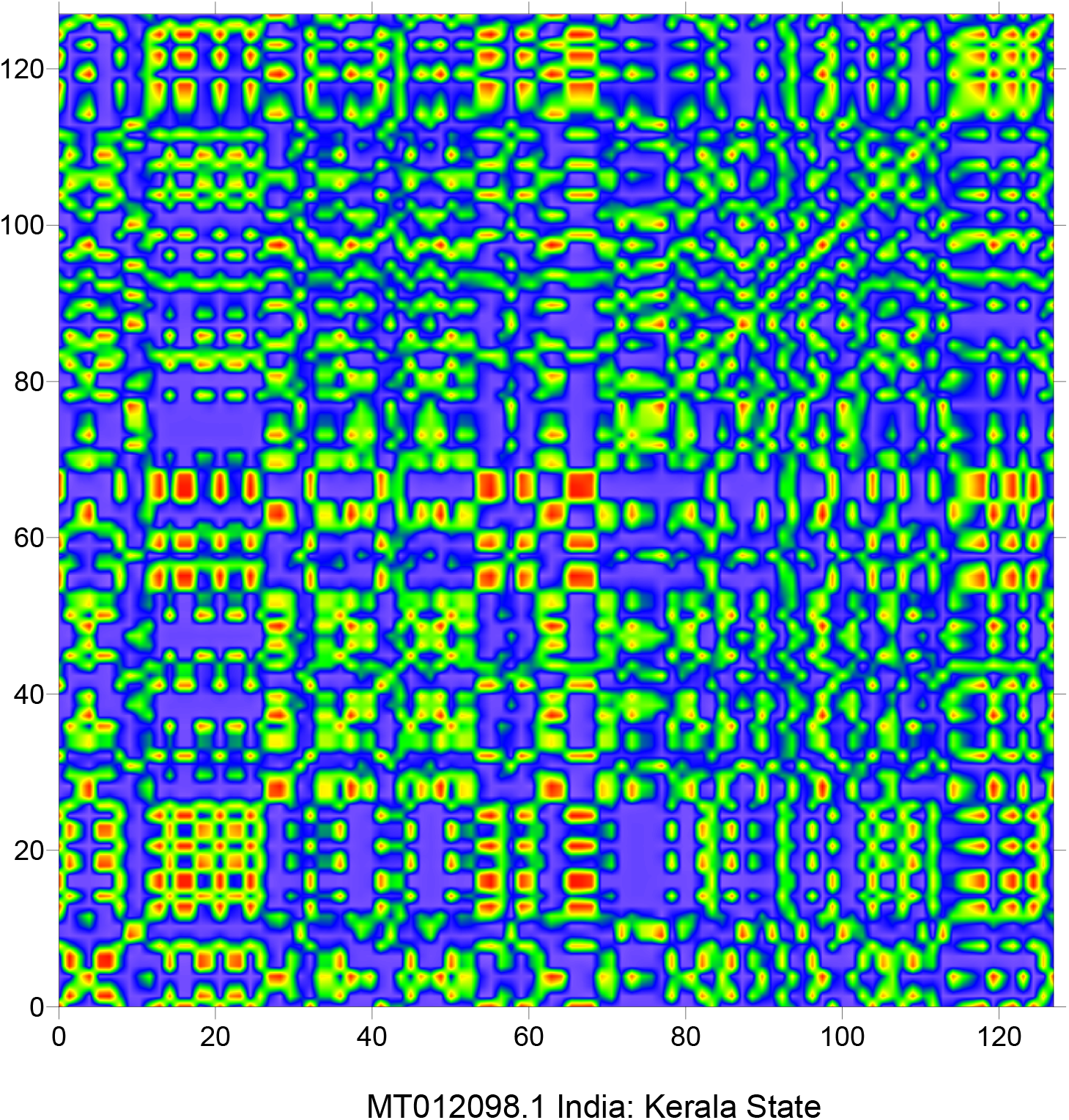
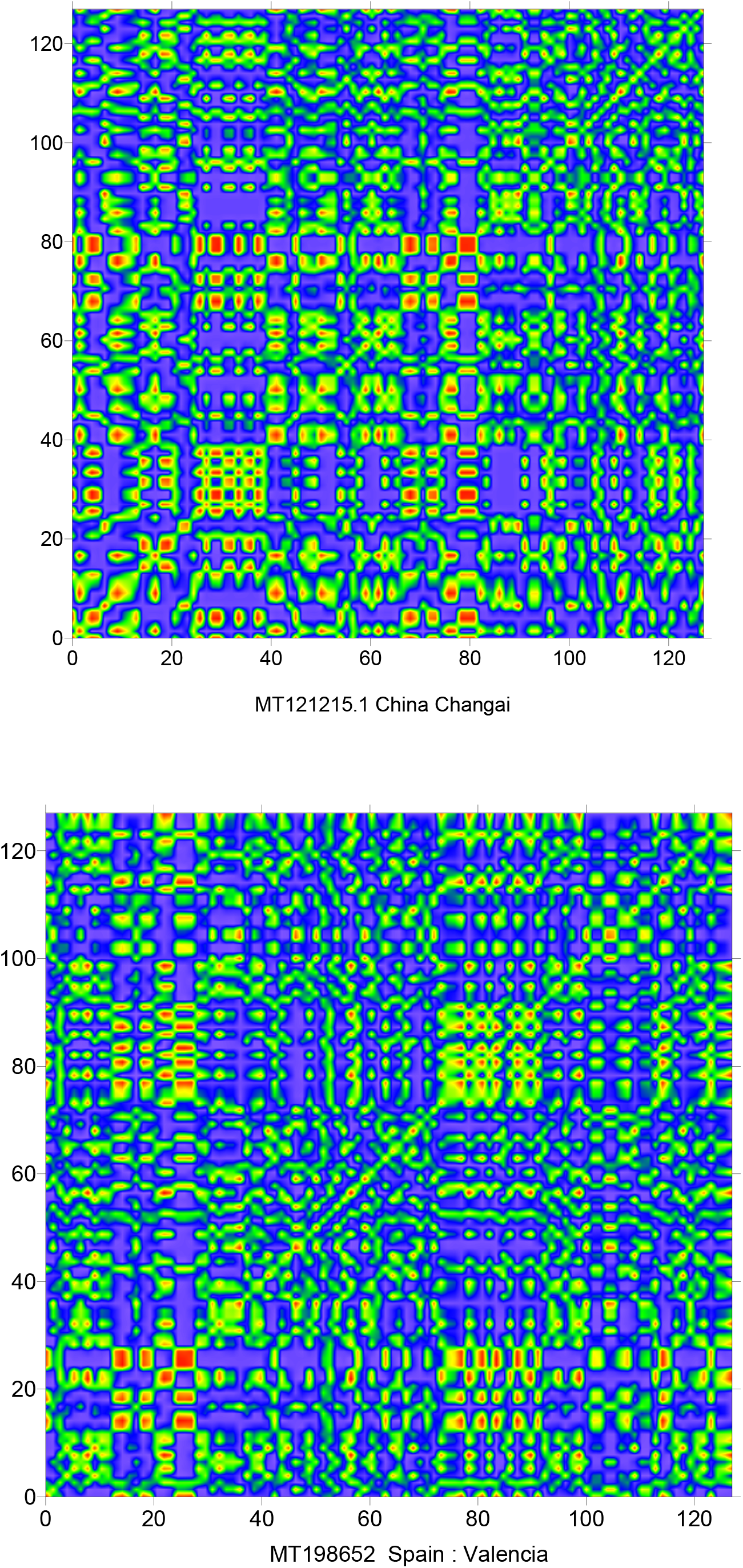
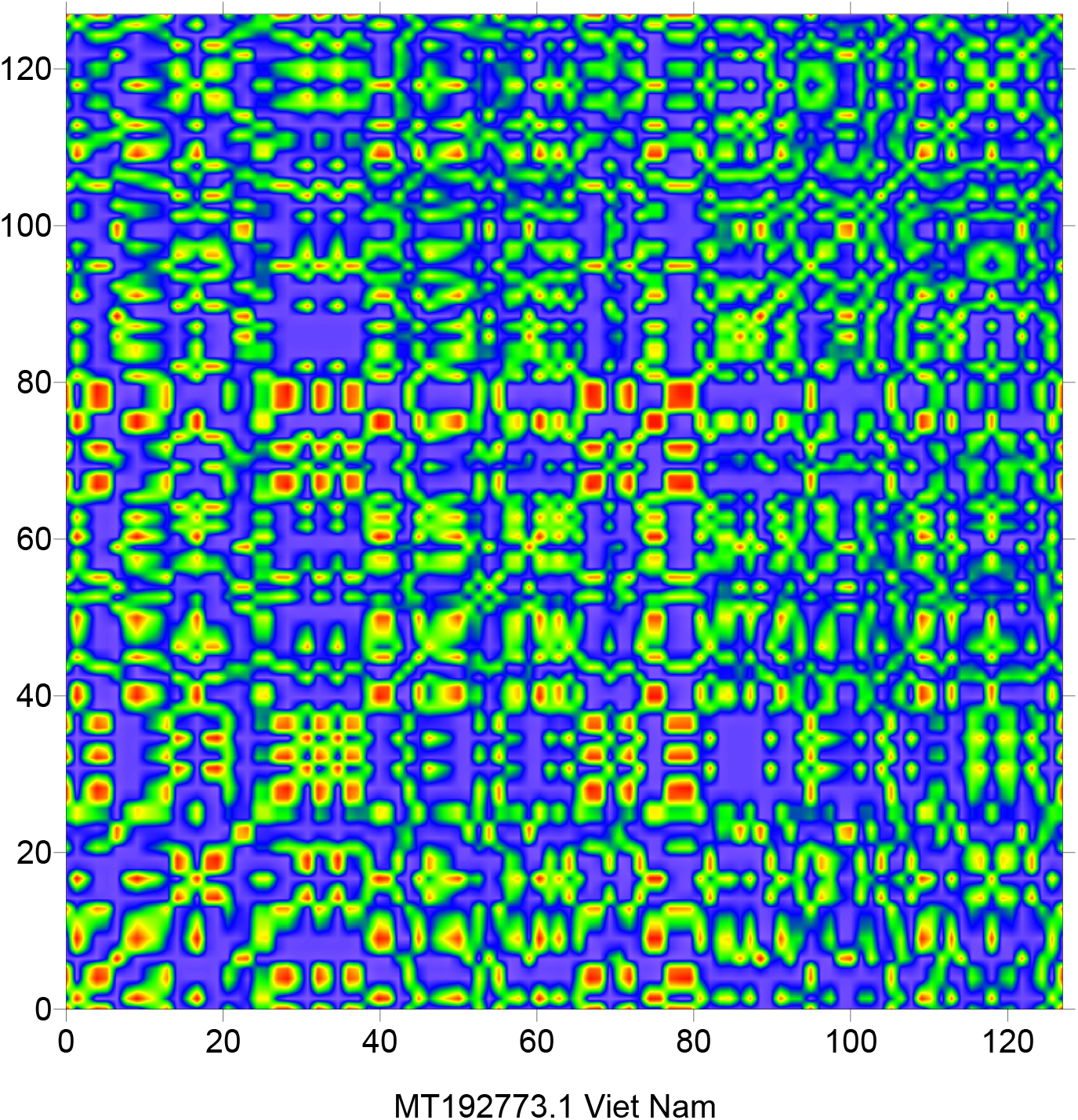
Indicator matrix for 21 sequences of the SARS-CoV2 coronavirus

## 5. Results discussion

The SARS-CoV2 coronavirus is characterized by a fractal dimension oscillating around 1.63, the obtained results are compared with SARS-CoV, SARS-like coronavirus and MERS-CoV. Figure 2 show the indicator matrix of a RNA sequence of SARS-CoV coronavirus (for more details about the SARS-CoV we invite the readers to the CDC report). We can observe that the patterns in this indicator matrix are similar to the patterns present in SARS-CoV2 indicator matrix, the fractal dimension of this sequence is equal to 1.60 which is similar to SARS-CoV2. Figure 3 shows the indicator matrix of 08 SARS-like coronavirus sequences (Conway, 2020), we can observe the patterns present in these maps are not similar to SARS-CoV2 coronavirus. The fractal dimension for each sequence is calculated (see table 03), these dimensions are oscillating around 1.58 (*D* = 1.58 ± 0.055), this value is slightly different to SARS-CoV2 fractal dimension. Figure 04 shows the indicator matrix of the MERS-CoV coronavirus (RNA sequence KT029139.1, downloaded from the NCBI database), it is clear that the patterns present here are similar to those of SARS-like coronavirus, the fractal dimension of MERS-CoV is 1.63 which is equal to the fractal dimension of SARS-CoV2 coronavirus. The directional wavelet transform signature of the four coronavirus are presented in figures 05, 06, 07 and 08, these signatures show a high similarity for the couples (SARS-CoV, SARS-CoV2) and (SARS-like, MERS-CoV).

**Figure 02:**
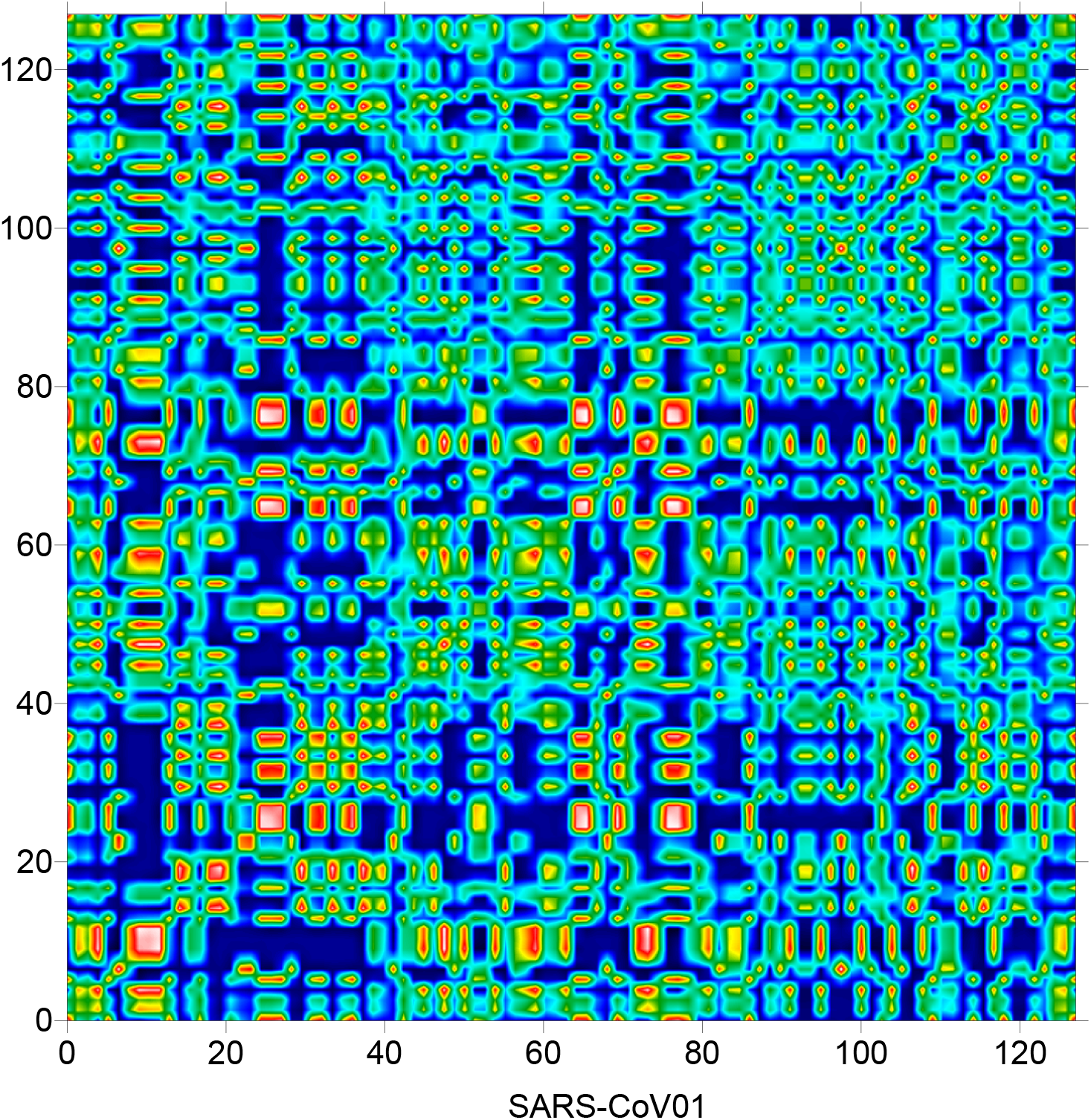
Indicator matrix of SARS-CoV coronavirus

**Figure 03:**
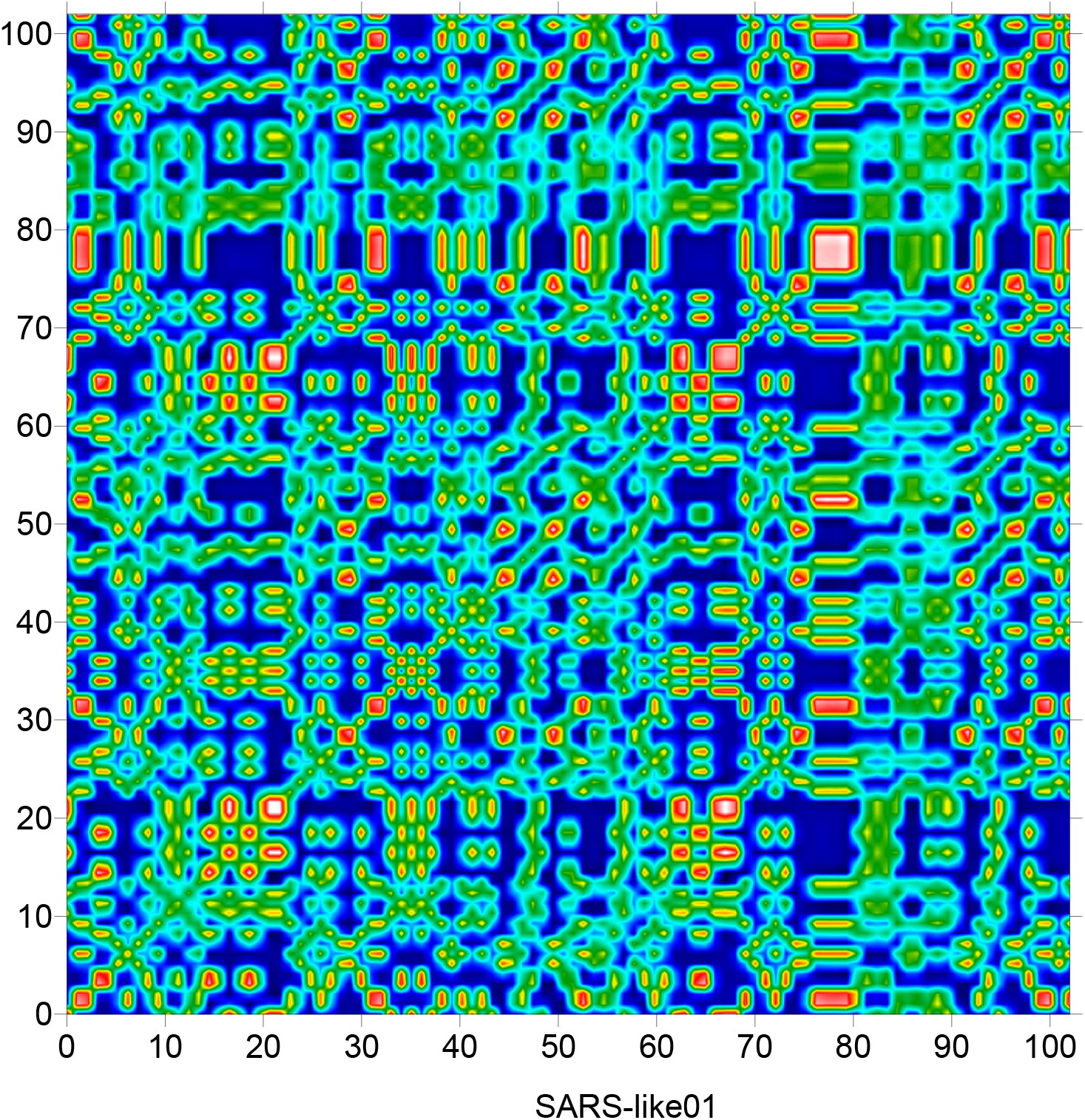
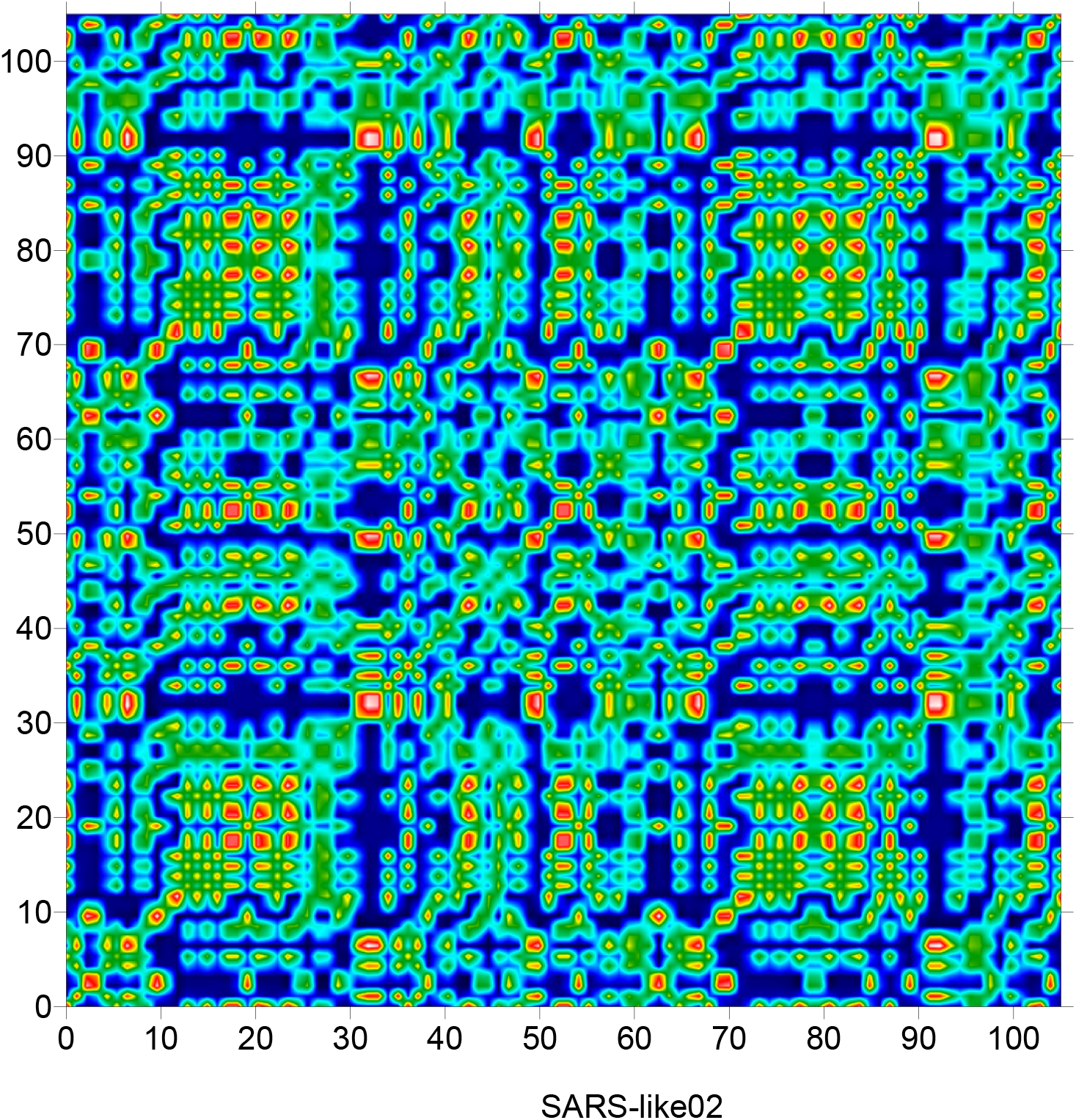
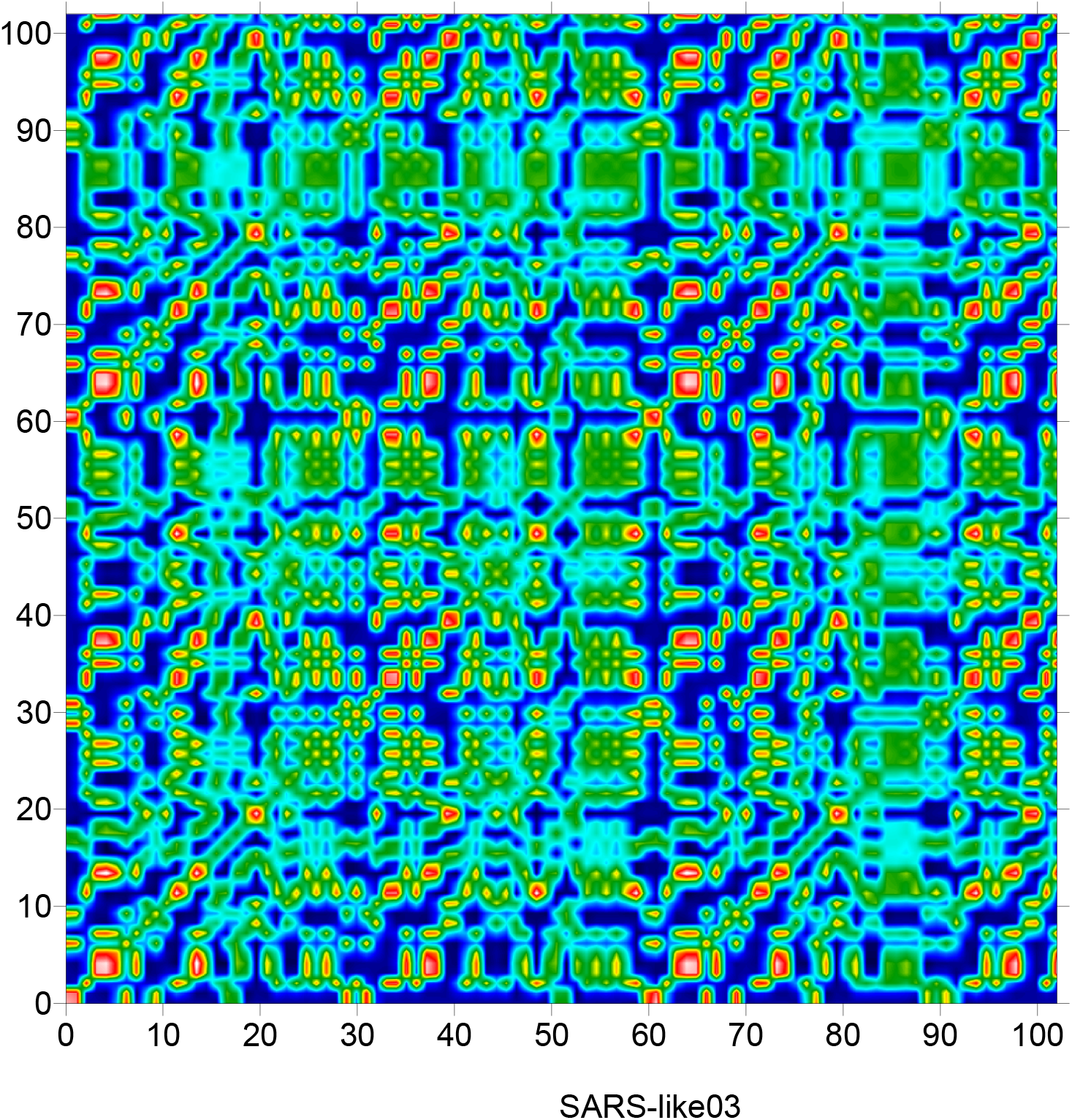
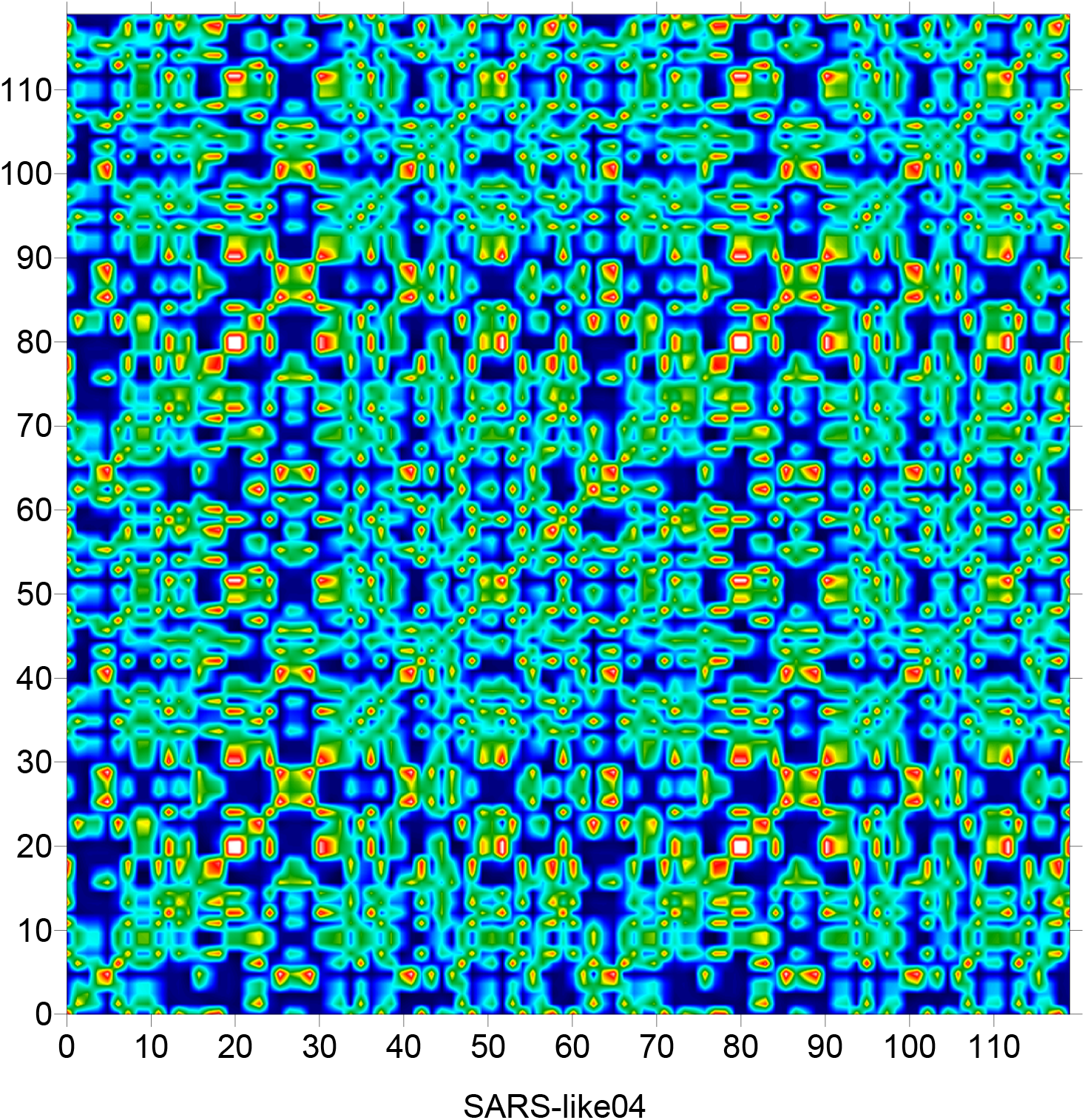
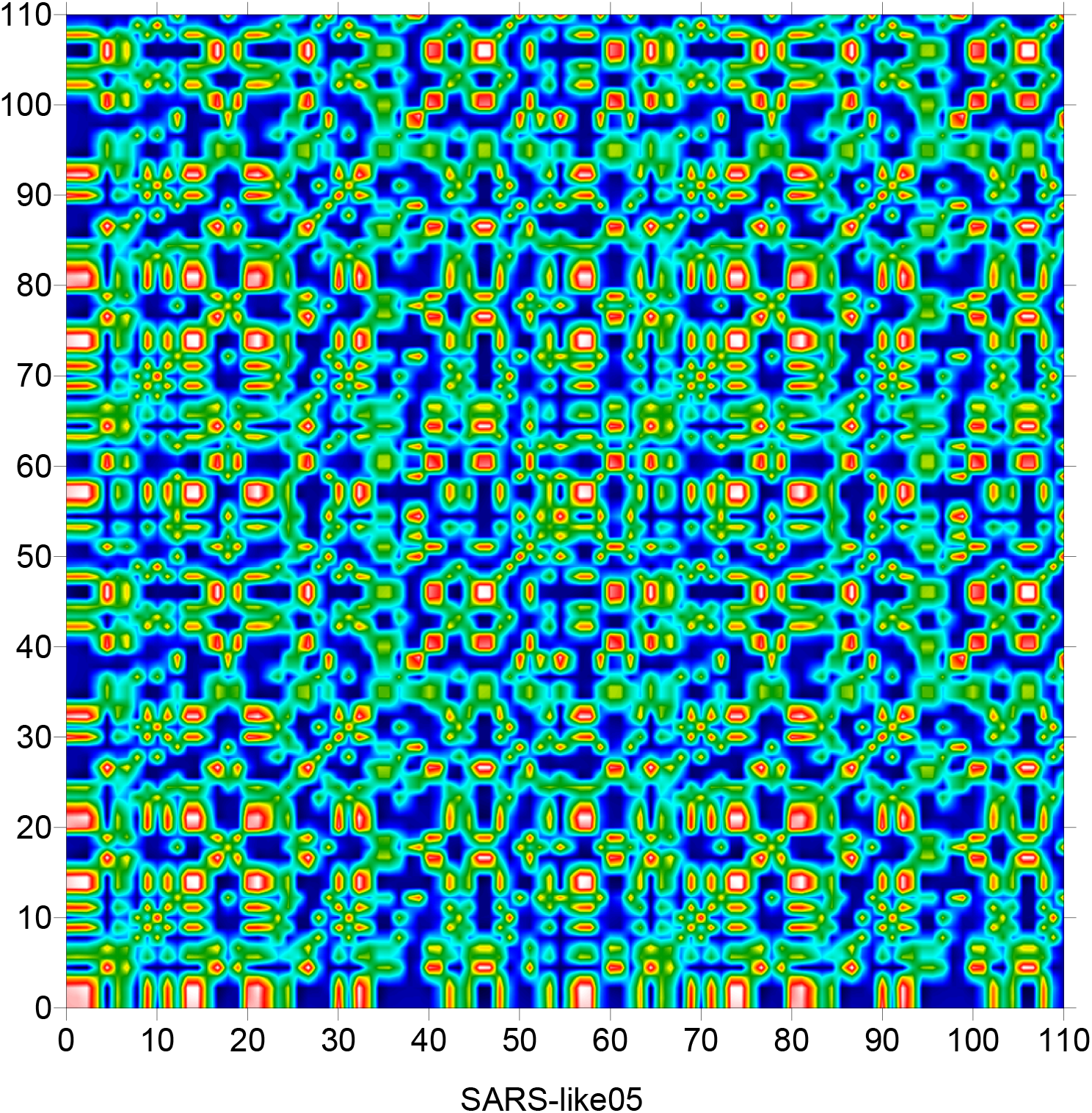
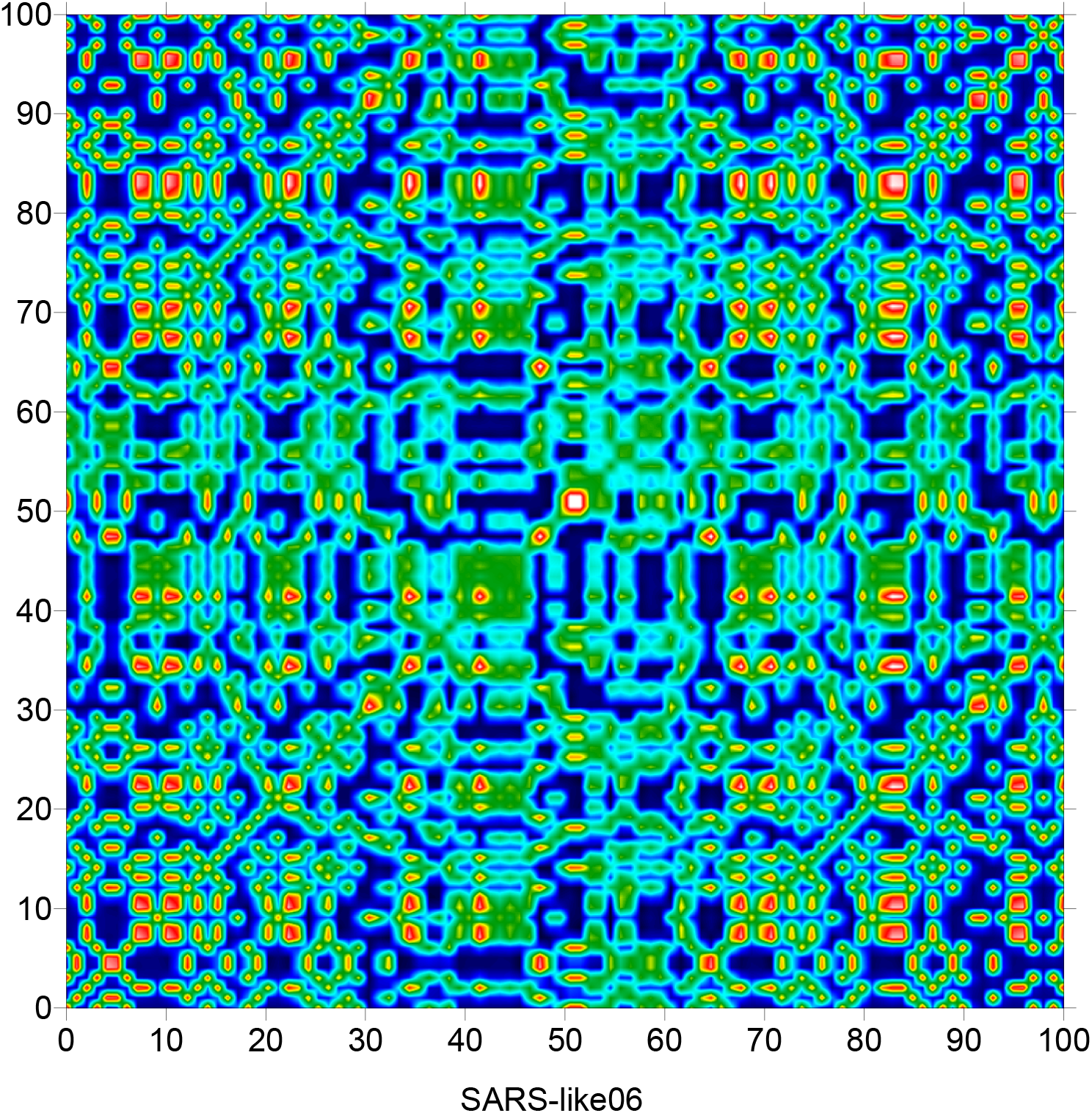
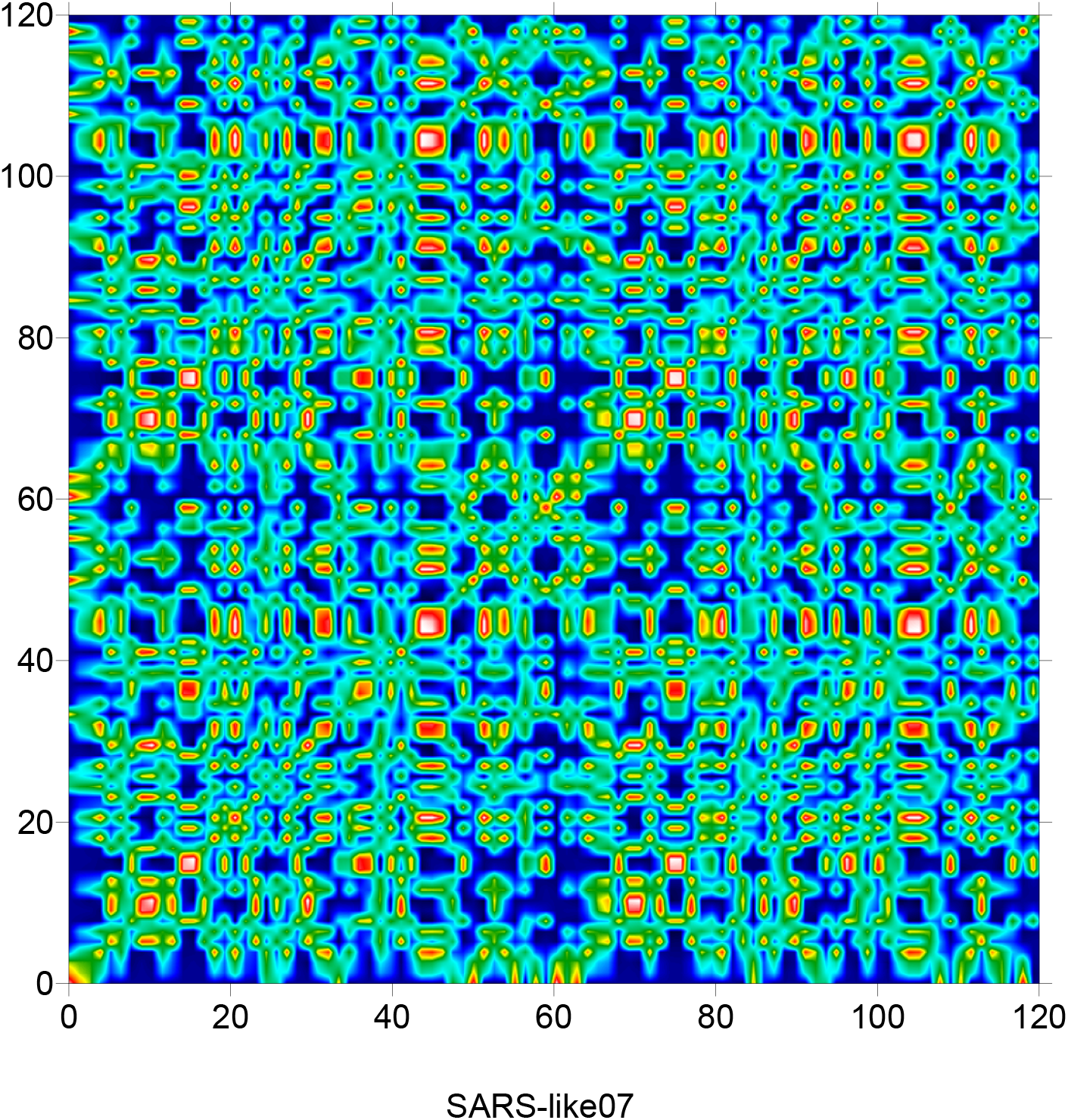
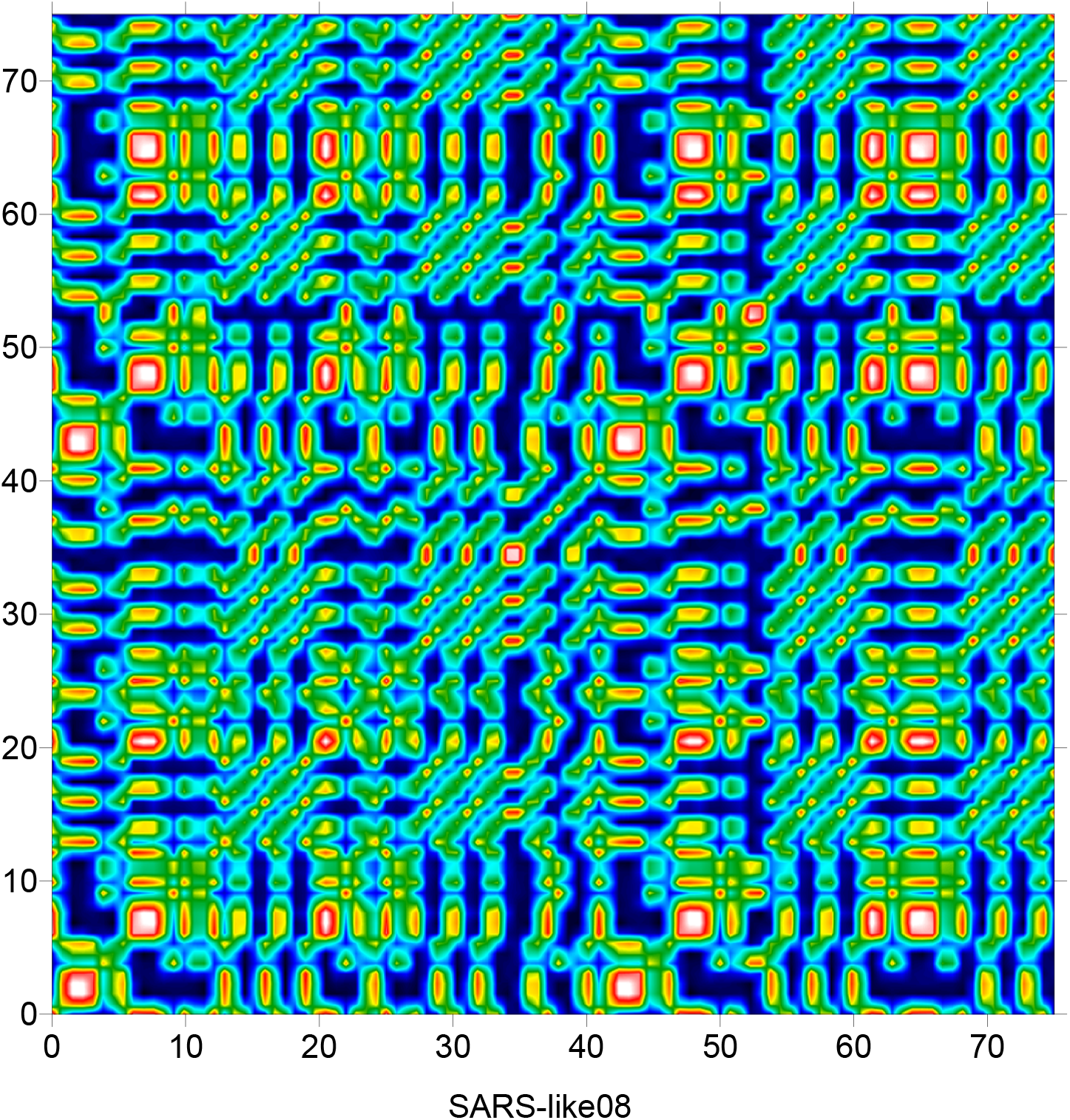
Indicator matrix of eight SARS-like coronavirus sequences

**Table 3.** Fractal dimension of 08 SARS-like coronavirus

|  | Fractal dimension |
| --- | --- |
| Seq-01 | 1.60 |
| Seq-02 | 1.59 |
| Seq-03 | 1.63 |
| Seq-04 | 1.58 |
| Seq-05 | 1.67 |
| Seq-06 | 1.59 |
| Seq-07 | 1.63 |
| Seq-08 | 1.60 |

**Figure 04:**
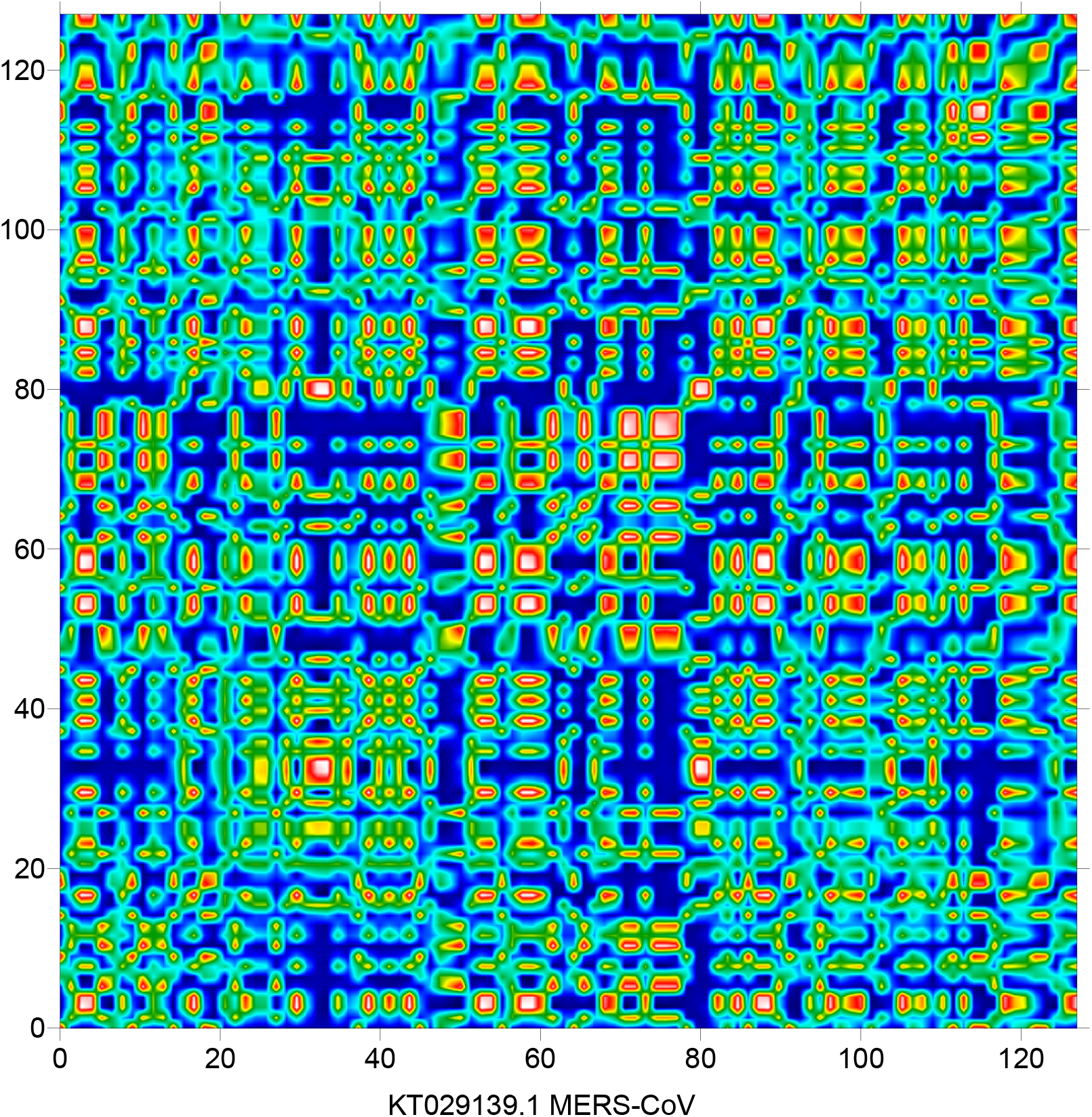
Indicator matrix of MESR-CoV coronavirus

**Figure 05:**
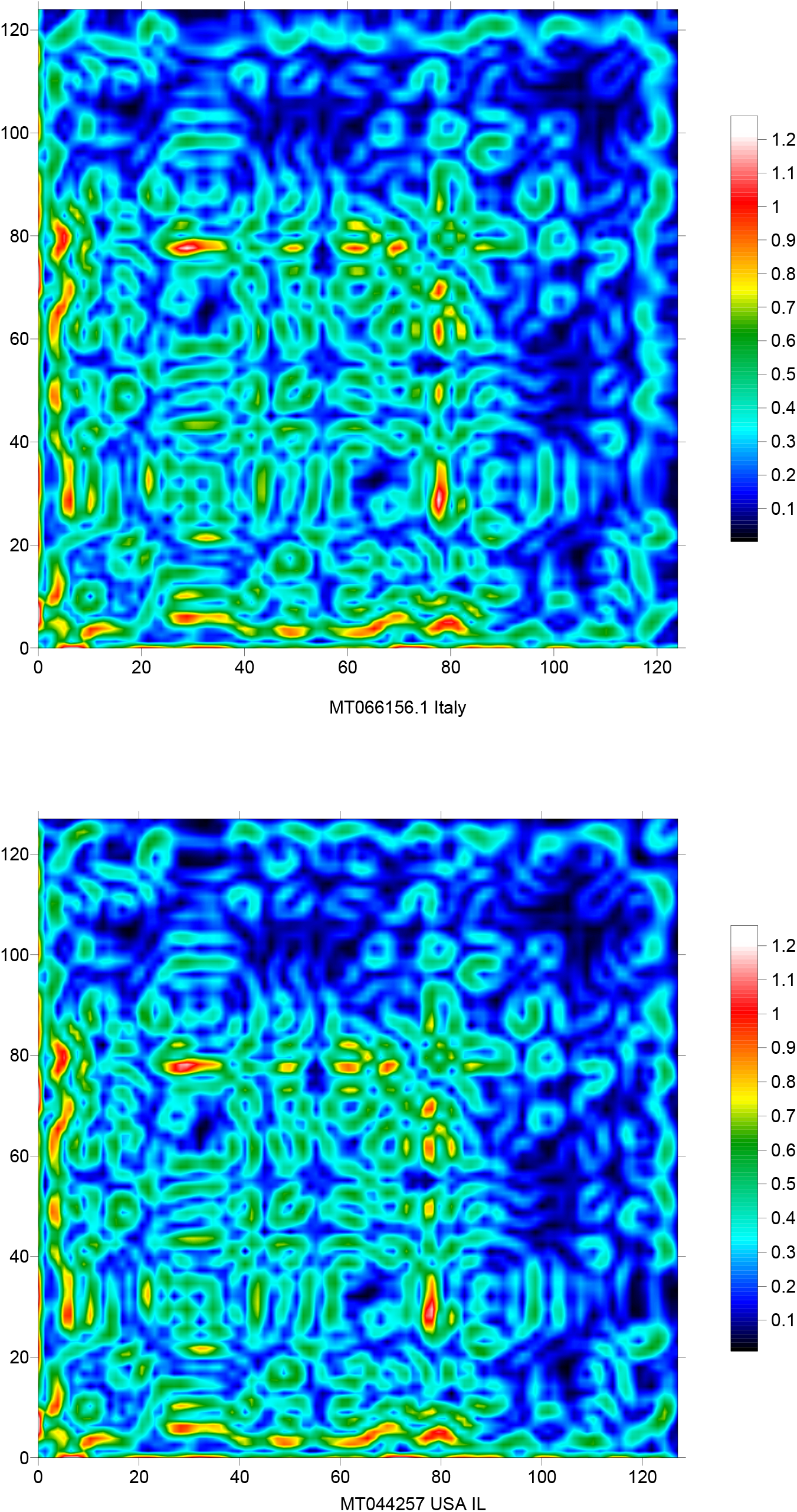
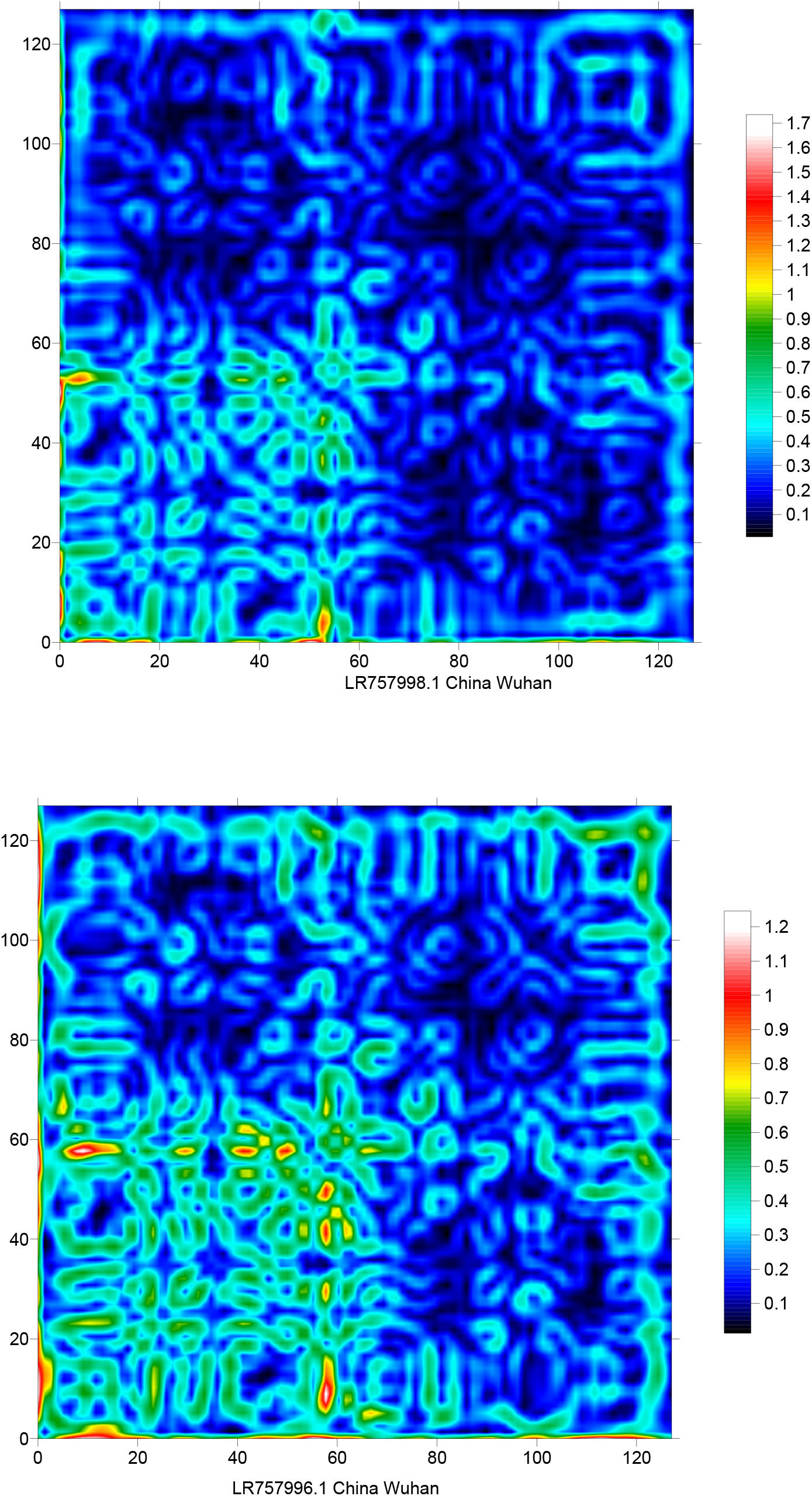
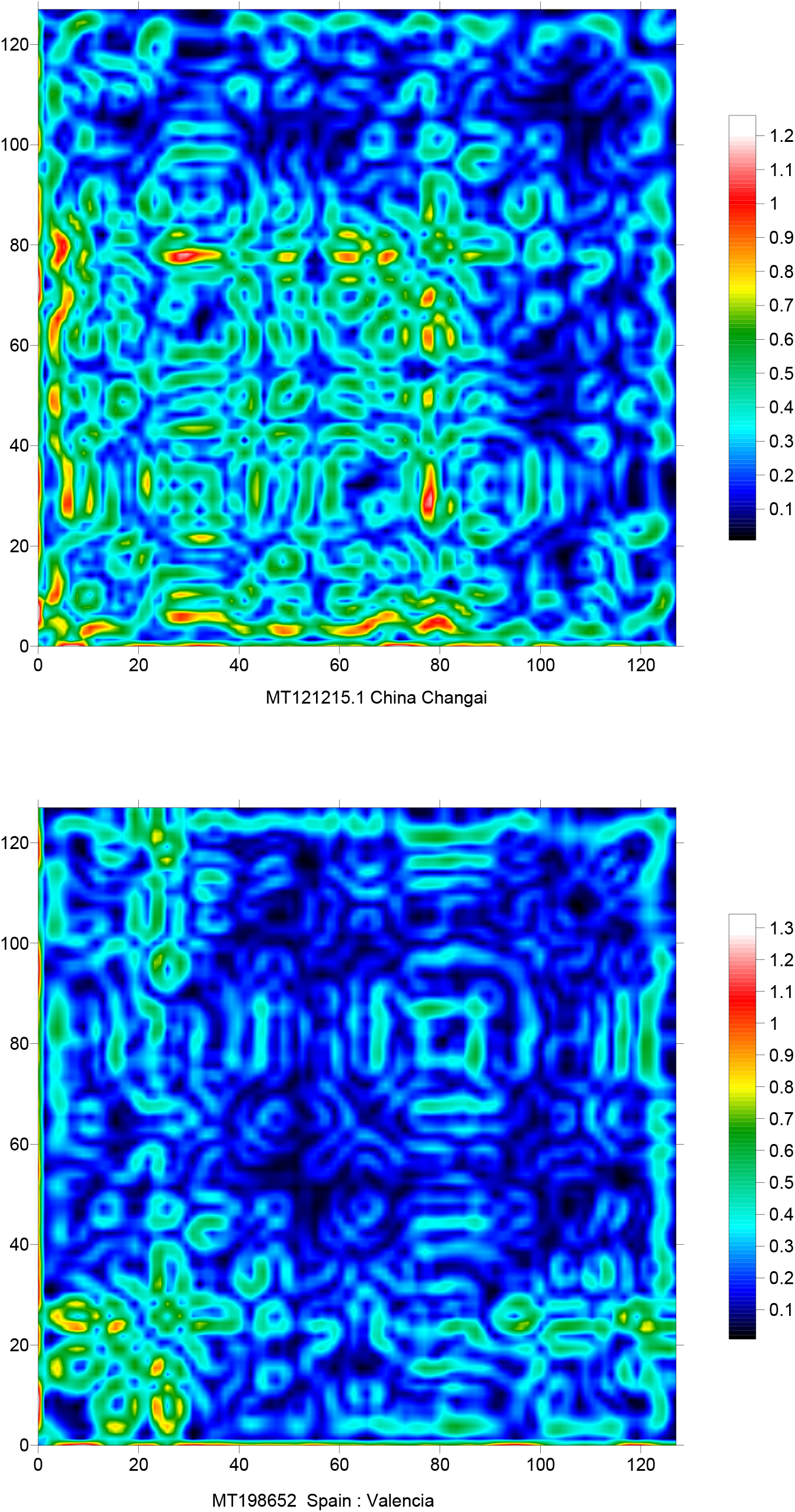
Directional wavelet transform signature of some SARS-CoV2 RNA sequences

**Figure 06:**
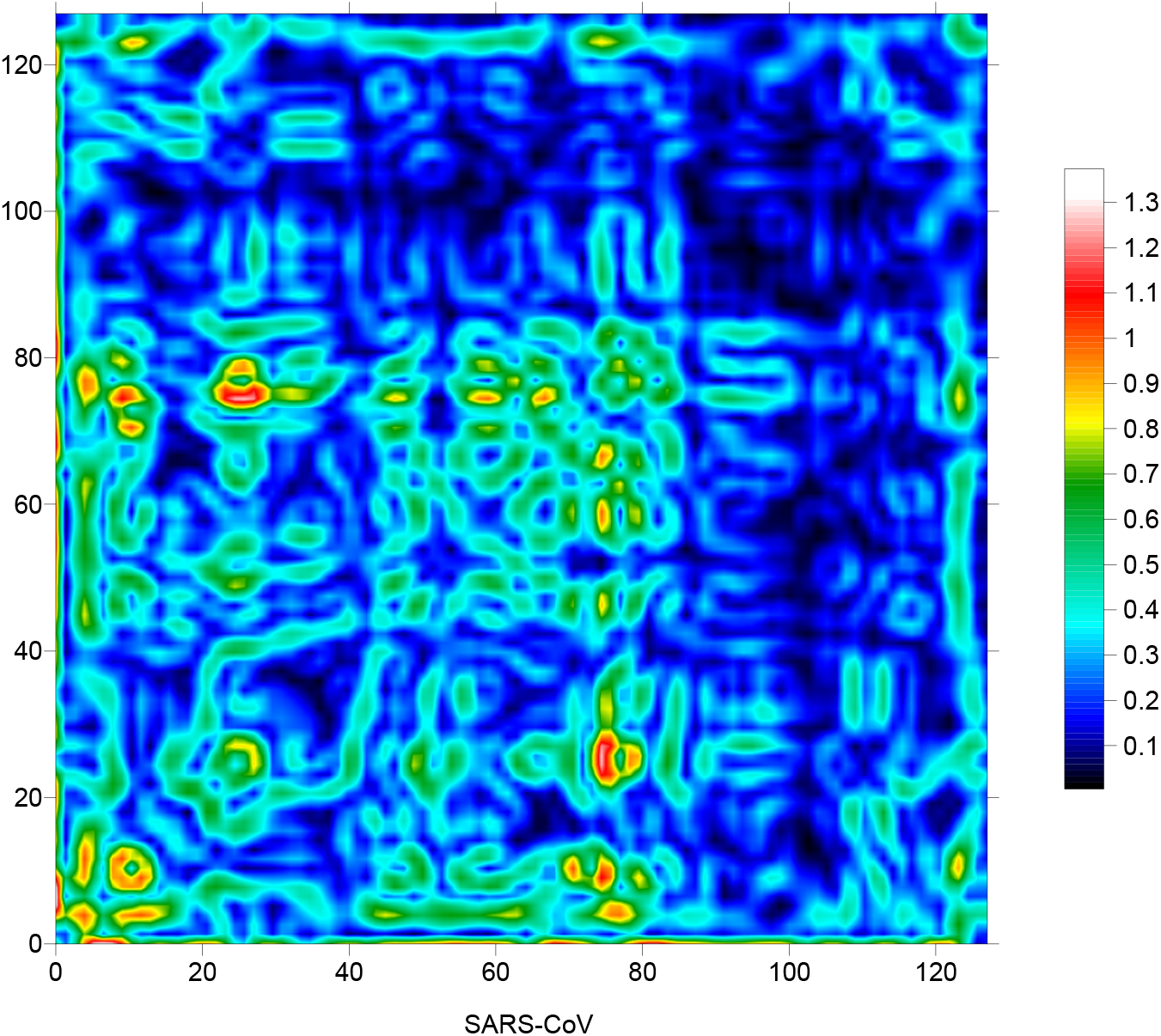
Directional wavelet transform signature of the SARS-CoV RNA sequence

**Figure 07:**
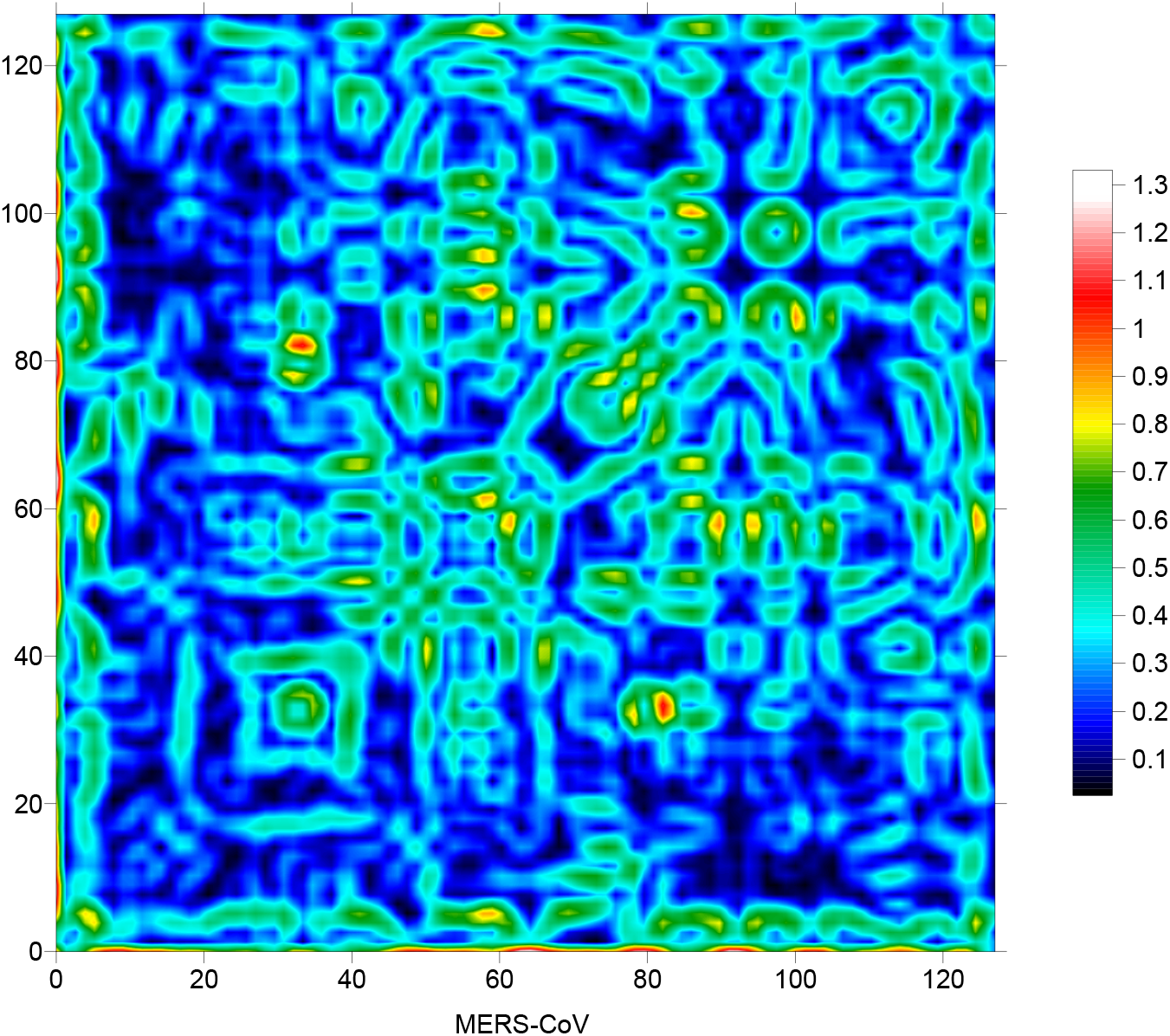
Directional wavelet transform signature of the MERS-CoV RNA sequence

**Figure 08:**
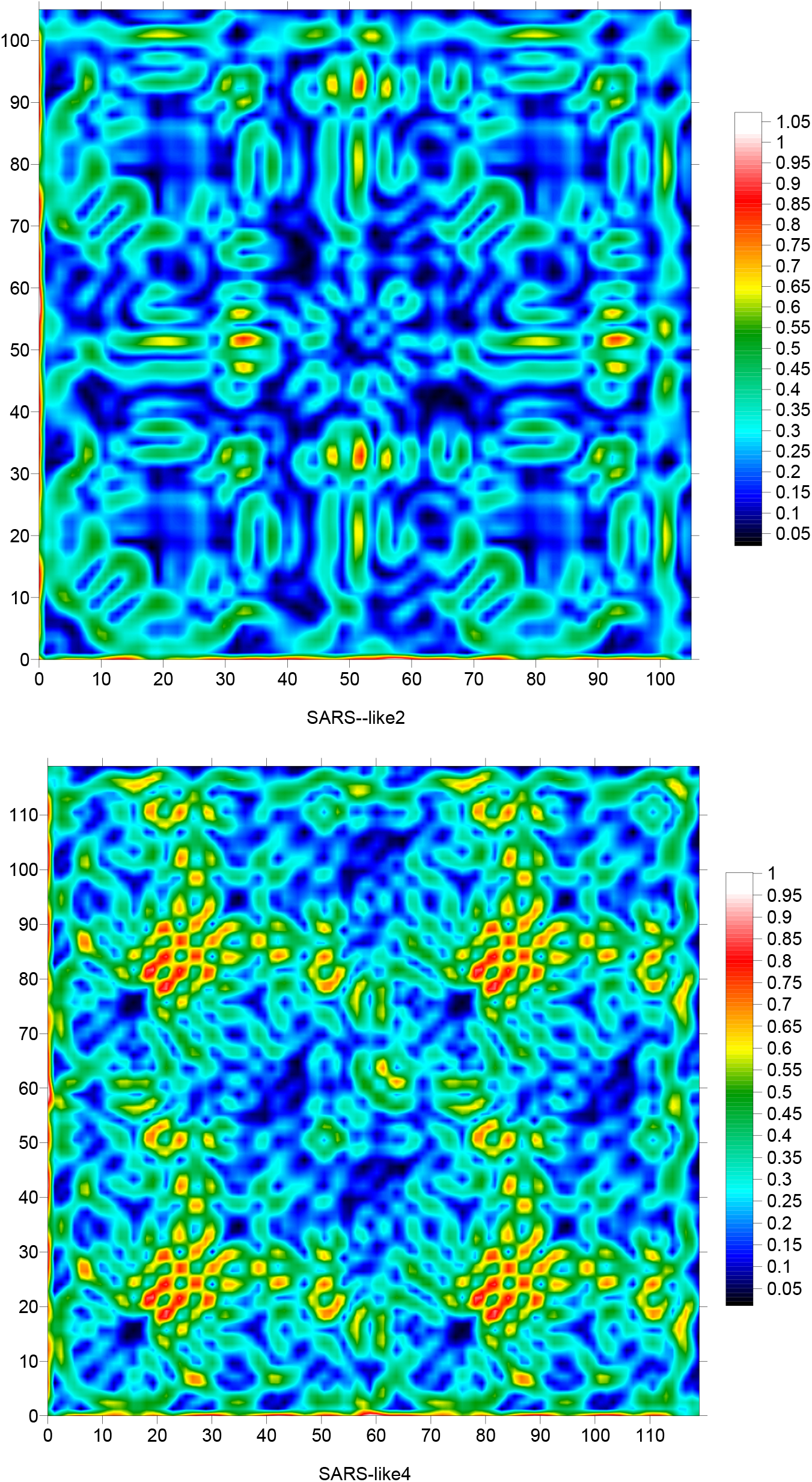
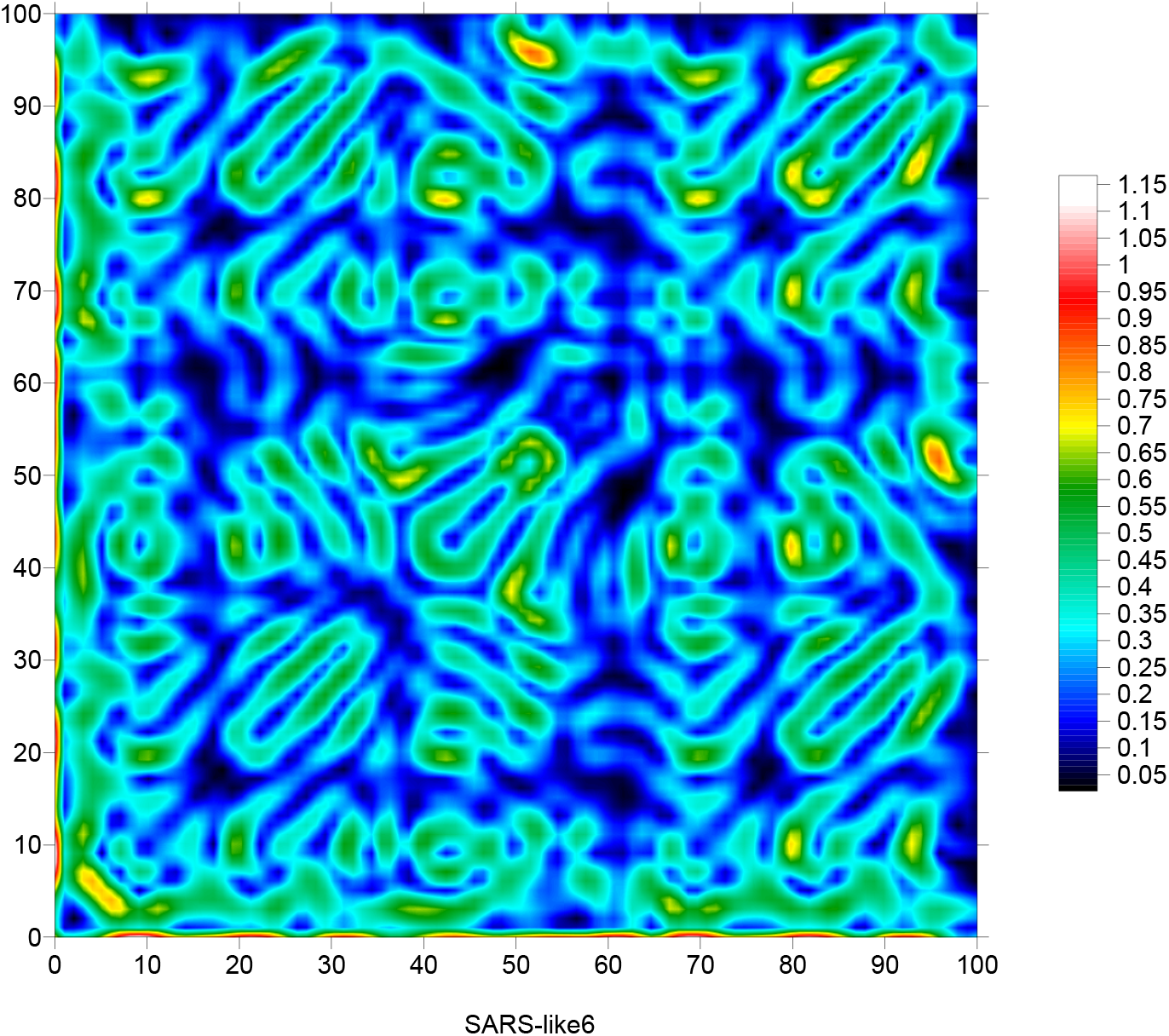
Directional wavelet transform signatures of four SARS-like coronavirus RNA sequences

## 6. Conclusion

We have analyzed 21 RNA sequences of SARS-CoV2 coronavirus, the indicator matrix and the fractal dimension of these sequences are calculated, obtained results are compared with those of SARS-CoV, MERS-CoV and SARS-like coronavirus, The fractal dimensions are identical or slightly different, however the maps of the indicator matrixes show the same patterns with different positions for the 21 RNA sequences of SARS-CoV2 with different positions. The 2D directional wavelet transform signatures show also the same patterns with different positions for these 21 RNA sequences. The indicator matrix and the 2D DCWT show similar patterns for (SARS-CoV2, SARS-CoV) and (MERS-CoV, SARS-like) coronavirus. By consequence the indicator matrix and 2D directional wavelet transform can be used to identify the presence of the coronavirus in human DNA genomes, these results enhance also the probability of SARS-CoV2 origin based on SARS-CoV mutation process.

